# Modification of amyloplast size in wheat endosperm through mutation of PARC6 affects starch granule morphology

**DOI:** 10.1101/2023.04.03.535339

**Authors:** Lara Esch, Qi Yang Ngai, J. Elaine Barclay, Rose McNelly, Sadiye Hayta, Mark A. Smedley, Alison M. Smith, David Seung

## Abstract

Starch granule morphology is a major factor determining the functional and nutritional properties of starch. Here, we reveal amyloplast structure plays an important role in starch granule morphogenesis in wheat endosperm. Wheat amyloplasts contain large discoid A-type granules and small spherical B-type granules. We isolated a mutant in durum wheat defective in the plastid division protein PARC6, which had increased plastid size in both leaves and endosperm. Endosperm amyloplasts of the mutant contained more A- and B-type granules than those of the wild type. In mature grains, the mutant had larger A- and B-type granules than the wild type, and its A-type granules had a highly aberrant, lobed surface. This defect in granule morphology was already evident at early stages of grain development, when granule size was identical between the mutant and the wild type, and occurred without obvious alterations in starch polymer structure and composition. Plant growth and photosynthetic efficiency, as well as the size, number and starch content of grains, were not affected in the *Ttparc6* mutants despite the large changes in plastid size. Interestingly, mutation of the PARC6 paralog, ARC6, in durum wheat did not increase plastid or starch granule size. We suggest this is because *Tt*PARC6 can complement disrupted *Tt*ARC6 function by interacting with PDV2, the outer plastid envelope protein that typically interacts with ARC6 to promote plastid division. We propose that amyloplast compartment size and available stromal volume play important roles in determining starch granule size, shape and number per amyloplast.

## Introduction

Amyloplasts are specialised non-green plastids, mostly found in roots, tubers and seeds, which synthesize starch (Sakamoto et al., 2008; Yun and Kawagoe, 2009; Jarvis and López-Juez, 2013; Sun et al., 2018). Starch is comprised of two glucose polymers, amylose and amylopectin, which together form semi-crystalline starch granules. In cereal grains, there is a large interspecies diversity in starch granule morphology and composition. This ranges from simple-type starch granules (e.g. in maize), stemming from a single initiation per amyloplast, to compound-type starch granules (e.g. in rice), where multiple starch granules are initiated simultaneously within one amyloplast (Matsushima et al., 2013; Chen et al., 2021). In wheat endosperm, there is a bimodal starch granule size distribution with large discoid A-type granules (18-20 µm diameter) and small spherical B-type granules (6-7 µm diameter) (Parker, 1985; Bechtel et al., 1990; Howard et al., 2011). Typically, one A-type granule is initiated in the amyloplast early during endosperm development. Smaller B-type granules are initiated later during endosperm development and at least partly within amyloplast stromules, thin tubular extensions of the plastid compartment, filled with stroma and surrounded by the plastid envelope (Parker, 1985; Bechtel et al., 1990; Langeveld et al., 2000; Howard et al., 2011; Hanson and Conklin, 2020). The broad range of starch granule size distributions found in cereal grains strongly influences the end use quality of starch (Chen et al., 2021).

Recently, advances in the understanding of starch granule initiation has enabled the identification of proteins that influence granule morphology in important staple crops like wheat and barley (Chia et al., 2020; Hawkins et al., 2021; Chen et al., 2022a). Given the occurrence of B-type granules in stromules, amyloplast morphology is also likely to play an important role in the spatial coordination of A- and B-type granule formation (Parker, 1985; Bechtel et al., 1990; Langeveld et al., 2000; Howard et al., 2011; Matsushima and Hisano, 2019). However, the role of amyloplast structure in determining starch morphology has not been studied in detail. Investigating structural components of amyloplasts in wheat could reveal important new insights on the formation of the unique bimodal granule morphology, and potentially provide new genetic targets for starch modification.

Plastid size and morphology are greatly influenced by plastid division. The division machinery consists of ring shaped protein complexes at the inner and outer envelope membranes [Filamenting temperature-sensitive mutant Z (FtsZ) and dynamin rings, respectively], which divide the plastids by binary fission (Miyagishima, 2011; Yoshida et al., 2012; Osteryoung and Pyke, 2014; Chen et al., 2018; Yoshida and Mogi, 2019). These contractile rings are coordinated by two sets of paralogous proteins that are important for transferring positional information from the FtsZ ring in the plastid stroma to the outside of the plastid, where the dynamin ring is formed (Chen et al., 2018; Yoshida and Mogi, 2019). Accumulation and Replication of Chloroplasts 6 (ARC6), a protein related to the cyanobacterial division protein Ftn2 (Vitha et al., 2003), spans the inner envelope membrane and tethers the FtsZ ring to the inner envelope membrane by interacting with its FtsZ2 subunits (Johnson et al., 2013). In the inter membrane space, the C-terminal domains of two ARC6 molecules interact with those of two Plastid Division 2 (PDV2) proteins to form a heterotetramer (Koksharova and Wolk, 2002; Vitha et al., 2003; Mazouni et al., 2004; Glynn et al., 2008; Marbouty et al., 2009; Wang et al., 2017). Paralog of ARC6 (PARC6) arose from an early duplication of ARC6 in vascular plants (Miyagishima et al., 2006; Glynn et al., 2009). It interacts with FtsZ2 in the chloroplast stroma and C-terminally with Plastid division 1 (PDV1), a protein probably also specific to vascular plants and originating from a duplication of PDV2 (Miyagishima et al., 2006; Glynn et al., 2009; Sun et al., 2023). PDV1 and PDV2 are responsible for recruitment of the Dnm2 (ARC5) subunits that form the outer dynamin ring at the plastid division site (Chen et al., 2018)

This model of plastid division by binary fission is mainly based on studies of Arabidopsis mesophyll chloroplasts (Chen et al., 2018), but there is evidence to suggest that the mechanism of division could differ between cell types, organs and species (Mingo-Castel et al., 1991; Bechtel and Wilson, 2003; Ishikawa et al., 2020). For example, Arabidopsis *parc6* mutants have giant chloroplasts in mesophyll cells (Glynn et al., 2009), but the effects of the *parc6* mutation on plastid morphology varies between different epidermal cell-types (Ishikawa et al., 2020): In pavement cells, the mutant has aberrant grape-like plastid morphology. In trichome cells, plastids exhibit extreme grape-like aggregations, without the production of giant plastids. Finally in guard cells, plastids are reduced in number, enlarged in size, and have activated stromules. Amyloplasts may also vary in their division mechanism. Dumbbell-shaped amyloplasts that appeared to undergo binary fission were observed in potato tubers (Mingo-Castel et al., 1991), but not in wheat endosperm, where it was proposed that amyloplasts rather divide through the formation of protrusions (Bechtel and Wilson, 2003). In rice endosperm, amyloplasts were shown to divide simultaneously at multiple sites, forming a beads-on-a-string like structure (Yun and Kawagoe, 2009).

Some of the conserved components of the plastid division apparatus appear to affect starch granule formation and morphology in amyloplasts (Tetlow and Emes, 2017). Disruption of *ARC6* in Arabidopsis greatly increased amyloplast size in root columella cells, and these appeared to contain larger starch granules (Robertson et al., 1995). Mutation of *ARC5* in rice resulted in either fused amyloplasts with thick connections or pleomorphic plastids with multiple division sites; and this was accompanied by an overall reduction in granulae size and irregular granule shape (Yun and Kawagoe, 2009). In potato, increased expression of *FtsZ1* resulted in fewer but larger starch granules within the tuber (De Pater et al., 2006). Whether these tubers had larger amyloplasts was not examined, but it is a possibility since in Arabidopsis, overexpression of *FtsZ* results in larger amyloplasts (Stokes et al., 2000). Due to the differences between species in starch granule initiation patterns in amyloplasts, it is difficult to predict the effect of altered amyloplast size on the initiation and morphogenesis of A- and B-type granules in wheat. As part of the broader goal to understand the relationship between amyloplast structure and starch granule biogenesis, we aimed to examine the role of ARC6 and PARC6 on plastid division in wheat endosperm and their impacts on A- and B- type starch granule initiation and morphogenesis.

## Results

### Identification of ARC6 and PARC6 genes in wheat

We first generated mutants in durum wheat (*Triticum turgidum* ssp. *durum*) defective in ARC6 and PARC6, to test whether they have increased amyloplast size in the endosperm. Using BLASTp against the bread wheat (*Triticum aestivum*) reference genome (Appels et al., 2018), we identified three putative homeologs of ARC6 encoded on group 6 chromosomes: *Ta*ARC6-A1 (TraesCS6A02G066200.3), *Ta*ARC6-B1 (TraesCS6B02G089500.2), *Ta*ARC6-D1 (TraesCSU02G117700.1) (Figs S1, S2a); as well as three putative homeologs of PARC6 encoded on group 2 chromosomes: *Ta*PARC6-A1 (TraesCS2A02G555400.1), *Ta*PARC6-B1 (TraesCS2B02G588400.1) and *Ta*PARC6-D1 (TraesCS2D02G559100.1) (Figs. S1, 1a). We confirmed these genes as orthologs of Arabidopsis ARC6 and PARC6 respectively, using phylogenetic tree analysis (Fig. S1). In the durum wheat (*Triticum turgidum* ssp. *durum*) reference genome (Svevo.v1, Maccaferri *et al*., 2019) the homeologs of *Ta*ARC6 corresponded to *Tt*ARC6-A1 (TRITD6Av1G015630.3) and *Tt*ARC6-B1 (TRITD6Bv1G022070.1), and their predicted amino acid sequences were identical to the bread wheat sequences. PARC6 in durum wheat corresponded to *Tt*PARC6-A1 (TRITD2Av1G286550.2) and *Tt*PARC6-B1 (TRITD2Bv1G255410.2). The predicted amino acid sequences from these primary gene models were 86.6% and 97.1% identical to their corresponding homeologs in bread wheat, respectively. Our analysis confirmed that both ARC6 and PARC6 genes are highly conserved in plants, and both durum and bread wheat have a single set of homeologs for each gene (Fig. S1).

### Phenotypic analysis Ttparc6 and Ttarc6 mutants

To isolate *Ttparc6* and *Ttarc6* mutants, we used the wheat TILLING mutant resource, featuring exome-capture sequenced, EMS-mutagenized mutants of durum cultivar Kronos (Krasileva *et al*., 2017). We obtained line Kronos1265 (K1265) that carries a premature stop codon in place of Gln503 in *TtPARC6-A1*, and Kronos2369 (K2369) carrying a premature stop codon in place of Gln456 in *TtPARC6-B1* (Fig. 1a). The K1265 and K2369 lines were crossed to create the *Ttparc6-1* and *Ttparc6-2* lines, arising from two independent crossing events using separate plants. KASP genotyping was used to identify homozygous single and double mutants for A- and B-genome mutations (*aa*BB, AA*bb*, *aabb*) and the corresponding ‘wild-type segregants’ (AABB) in the F2 and F3 generation. We also crossed the *Ttparc6-2* double mutant with a transgenic amyloplast reporter line in cultivar Kronos, to serve two purposes: first, this transgenic line was not exposed to EMS mutagenesis and was therefore a suitable genetic background for backcrossing to remove undesirable background mutations in *Ttparc6-2*. Second, the line carries a single transgene encoding an mCherry protein targeted to the plastid stroma, enabling visualisation of amyloplasts (Matsushima and Hisano, 2019). KASP and PCR-based genotyping were used to isolate backcrossed (BC) individuals for each *PARC6* genotype (BC AABB, BC *aa*BB, BC AA*bb,* BC *aabb*); as well as double mutant and wild-type segregant lines carrying the reporter transgene (*Ttparc6-2* + *cTPmCherry aabb* and *Ttparc6-2* + *cTPmCherry* AABB).

Under our growth conditions, the single or double mutant lines for *Ttparc6* were identical to their wild-type controls with respect to growth, development and number of tillers (Fig. 1b, c Fig. S3a, b, l). To examine whether the mutations affected chloroplast size in leaves, we used confocal microscopy on mesophyll cells isolated from the youngest fully developed leaf of seedlings. Chloroplast size was drastically increased in both backcrossed and non-backcrossed double mutant lines compared to their wild-type controls (Fig. 1d-h). In the single mutants (*Ttparc6-1* AA*bb*, *Ttparc6-1 aa*BB, *Ttparc6* BC AA*bb* and *Ttparc6* BC *aa*BB), chloroplast size was visually indistinguishable from the wild-type controls (Fig. S3c-j).

**Figure 1:**
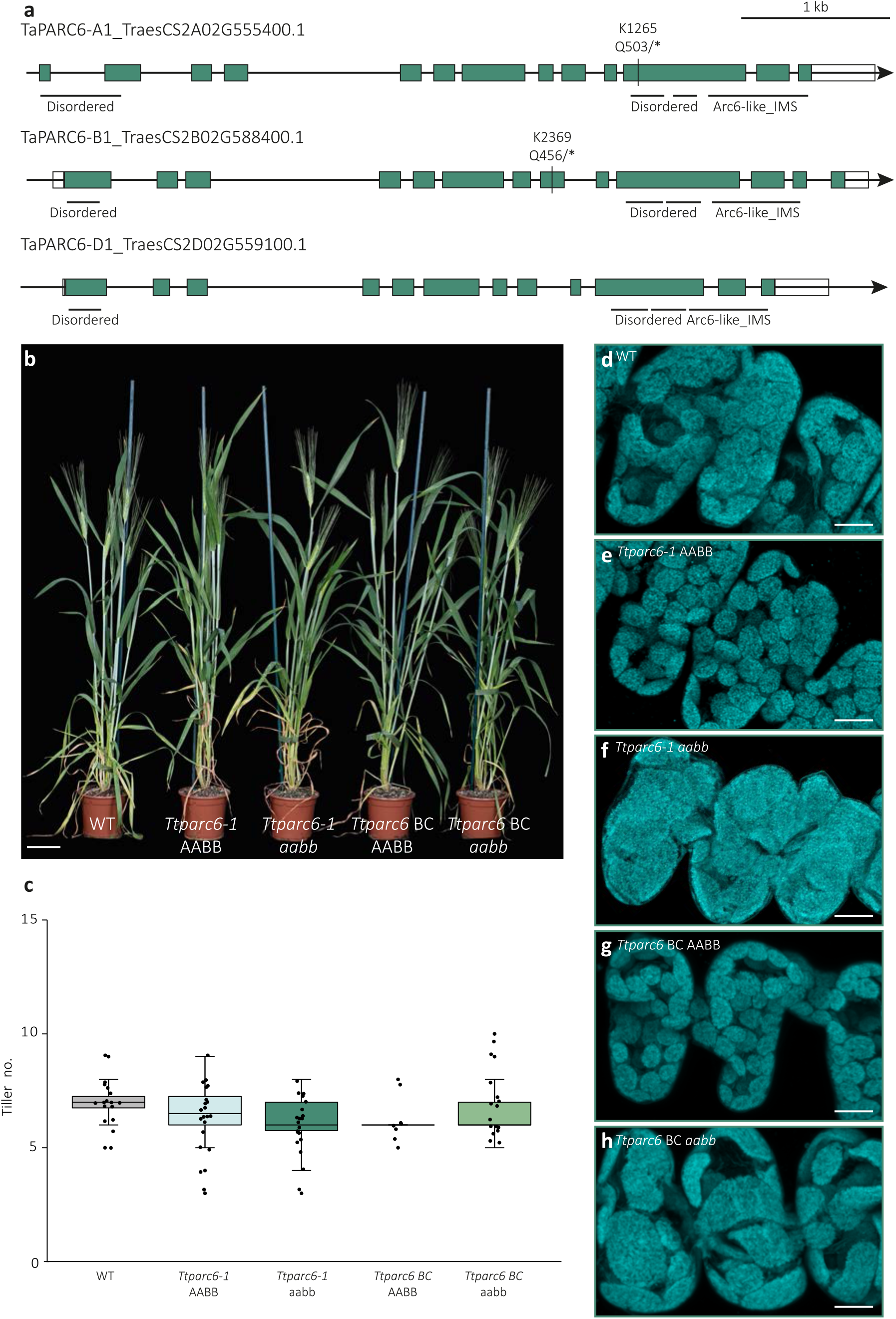
Growth phenotype of *Ttparc6* mutants. **(a)** Schematic illustration of the gene models for the primary transcripts of *TaPARC6-A1*, *-B1 and -D1* in bread wheat. Exons are represented as teal boxes and UTRs are represented as white boxes. Mutation sites in K1265 and K2369 are indicated by black lines and the resulting amino acid to stop codon (*) substitutions are annotated. Regions encoding domains are indicated by black horizontal lines (IMS: Inter Membrane Space). **(b)** Photograph of 8-week old *Ttparc6* TILLING double mutant (*Ttparc6-1 aabb*), the corresponding wild-type segregant (*Ttparc6-1 AAB*B), *Ttparc6* backcrossed double mutant (*Ttparc6 BC aabb*), the corresponding wild-type segregant (*Ttparc6 BC AABB*) and WT wheat (cv. Kronos) plants. Bar = 10 cm. **(c)** The number of tillers per plant (Tiller no.) of mature *Ttparc6* mutant plants. The top and the bottom of the box represent the lower and upper quartiles respectively, and the band inside the box represents the median. The ends of the whiskers represent values within 1.5x of the interquartile range. Outliers are values outside 1.5x the interquartile range. There is no significant difference between the lines as determined by Kruskal-Wallis one-way ANOVA on ranks (p = 0.097). **(d-h)** Images of mesophyll-cell chloroplasts in the third leaf of *Ttparc6* mutant seedlings. Images were acquired using confocal microscopy and are Z-projections of image stacks. Chlorophyll auto-fluorescence of the chloroplasts is shown in cyan. Bar = 10 μm.

Since the *Ttparc6* double mutants had increased chloroplast size but seemingly normal growth, we examined their photosynthetic efficiency using gas-exchange analysis (Fig. S4). In light response curves, both the backcrossed and non-backcrossed double mutants showed a slight tendency towards decreased photosynthesis rates (A) compared to the wild-type controls (Fig S4a-e), but statistical analysis of extracted A values at ambient (280 µmol m^-2^ s^-1^) or high light (2000 µmol m^-2^ s^-1^) revealed no differences between the mutants and wild-type controls (Fig. S4f). There were also no significant differences between the mutants and wild-type controls in maximal carboxylation rate (Vcmax) (Fig S4g). Although the maximal electron transport (Jmax) was significantly decreased in the backcrossed double mutant (*Ttparc6* BC *aabb*) compared to its backcrossed wild-type segregant (*Ttparc6* BC AABB), it was not significant when compared to the wild type. The Jmax of the non-backcrossed *Ttparc6-1 aabb* double mutant was also not different to its wild-type controls (Fig S4h). Therefore, we did not detect any consistent effect of the *Ttparc6* mutations on the measured photosynthetic parameters.

Using an approach similar to that for *TtPARC6*, we also isolated a mutant defective in both homeologs of *TtARC6*, crossing the lines Kronos3404 (K3404) and Kronos2205 (K2205) to introduce premature stop codons in place of Gln631 in *TtARC6-A1* and in place of Trp647 in *TtARC6-B1* (Fig. S2a). Unlike both wheat *Ttparc6* mutants described above and Arabidopsis *arc6* mutants (Vitha et al., 2003), chloroplast size was unaffected by the *Ttarc6* mutations (Fig. S2b, c). Therefore, we focused our analyses of endosperm starch on the *Ttparc6* mutants.

### Ttparc6 mutants have normal grain size, number and yield

We examined the grains harvested from the *Ttparc6* mutants (Fig. 2a, b, Fig. S3k). The total grain yield per plant for the backcrossed and non-backcrossed double mutants, as well as for the single mutants, was not significantly different from their wild-type controls (Fig. 2c, S3m). There was also no consistent effect of *Ttparc6* mutations on grain size, weight, and total starch content (Fig. 2d-, S3o, p). The average thousand grain weight (TGW) for the non-backcrossed *Ttparc6* double and single mutants was not significantly different to the wild-type controls (Fig. 2d, S3n). However, the TGW of the backcrossed *Ttparc6 BC aabb* double mutant was significantly higher than in the wild-type controls (24% increase relative to the WT) (Fig. 2d). Grain size parameters were also significantly higher in this backcrossed double mutant than in the wild-type segregant and the WT (area, width and length increased by 15%, 10% and 6%, respectively) (Fig 2f-h). Since these increases in grain weight and size were only observed in the backcrossed double mutants and not in the non-backcrossed double mutant, they are unlikely to be directly caused by the *Ttparc6* mutations.

**Figure 2:**
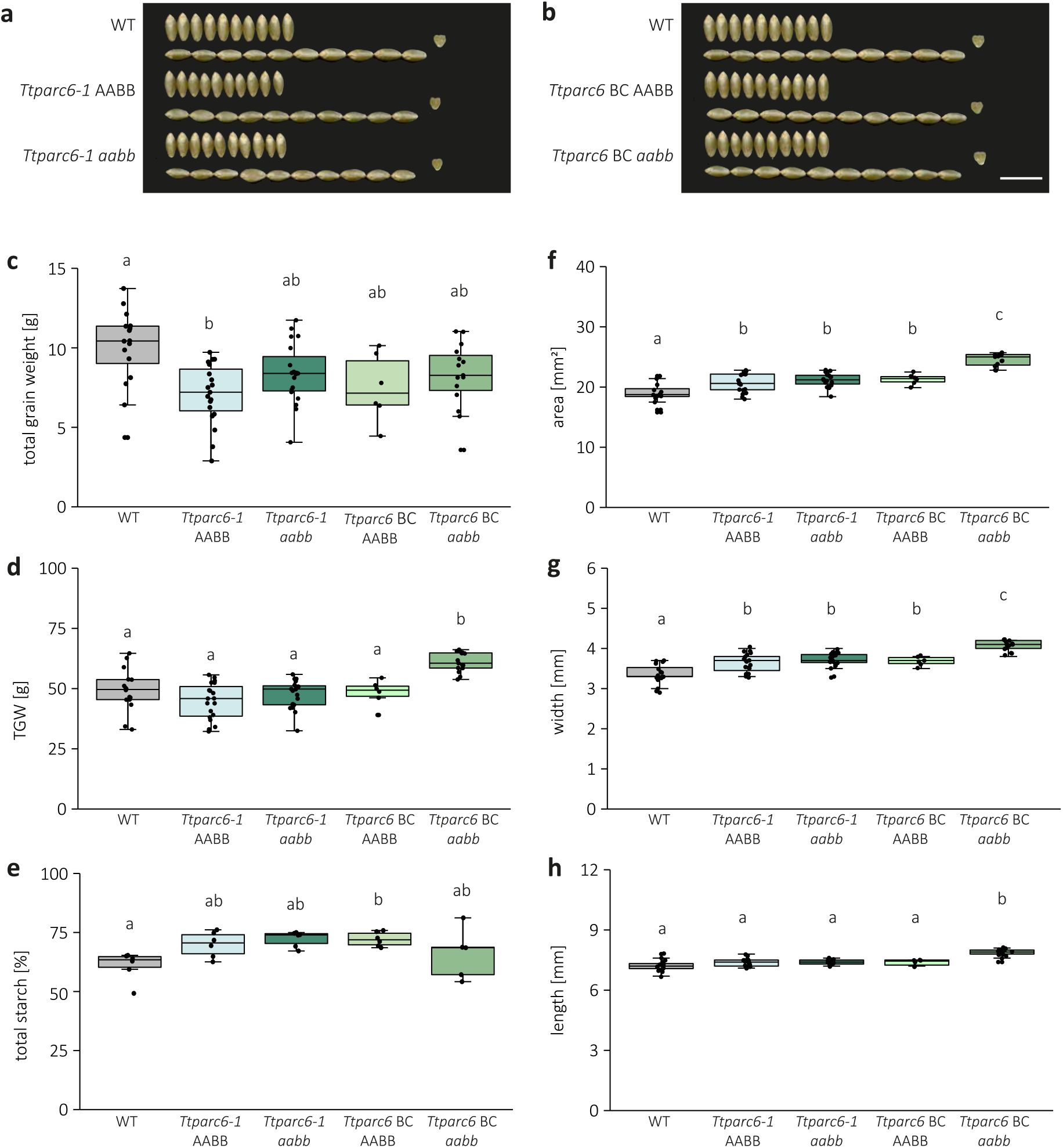
Seed phenotype of *Ttparc6* mutants. **(a&b)** Photographs of ten representative mature grains per genotype. Note that the same WT grains were used in both panels. Bar = 1 cm. **(c)** Total grain weight harvested per plant (in g). Dots represent the total grain weight of individual plants (*n* = 6-19) per genotype. Significant differences under a one-way ANOVA and all pairwise multiple comparison procedures (Tukey’s test) are indicated with different letters (p ≤ 0.002). **(d)** Thousand grain weight (TGW) (in g). Dots represent calculated TGW of individual plants (*n* = 6-19) per genotype. Significant differences under a one-way ANOVA and all pairwise multiple comparison procedures (Tukey’s test) are indicated with different letters (p ≤ 0.001). **(e)** Total starch content as % (w/w). Three technical replicates of two biological replicates per genotype. Significant difference between the genotypes under a Kruskal-Wallis one-way ANOVA on ranks and all pairwise multiple comparison procedures (Tukey’s test) are indicated with different letters (p ≤ 0.041). **(f-h)** Grain size parameters measured as seed area (f), width (g) and length (h). Dots represent the average for each parameter calculated from grains from individual plants (*n* = 6-19) per genotype. Significant differences under a one-way ANOVA (for f and g) or a Kruskal-Wallis one-way ANOVA on ranks (for h), and all pairwise multiple comparison procedures (Tukey’s test) are indicated with different letters (p ≤ 0.001). For all boxplots, the bottom and top of the box represent the lower and upper quartiles respectively, and the band inside the box represents the median. The ends of the whiskers represent values within 1.5x of the interquartile range, whereas values outside are outliers.

### Ttparc6 mutants have increased starch granule sizes in the endosperm

We purified starch granules from mature grains of the *Ttparc6* mutants and examined starch granule size and morphology. Using a Coulter counter, we observed that all genotypes had a bimodal distribution of starch granule size (Fig. 3a, b). In the wild type, the A-type granule peak had its maximum at around 19 µm diameter and the B-type granule peak maximum was at about 6 µm diameter. The *Ttparc6* double mutants had drastically altered granule size distributions compared to the WT and the corresponding wild-type segregants. The A-type granule peak in the mutants was shifted towards larger granule diameters; the B-type granule peak was not only shifted to larger granule sizes but also had a larger peak area. We fitted a log-normal distribution to the B-type granule peak and a normal distribution to the A-type granule peak to derive the mean diameters of A- and B-type granules, as well as the B-type granule content (percentage of total starch volume that is present as B-type granules). The mean diameter of A-type granules of both double mutant (*aabb*) genotypes were significantly larger than those of their corresponding wild-type controls (15.1% increase in *Ttparc6-1 aabb* and 21.9% increase in *Ttparc6 BC aabb*) (Fig. 3c). While there was a shift towards larger B-type granule sizes in both the *Ttparc6-1 aabb* double mutants, only the backcrossed double mutant genotype had a significant increase in the mean diameter of B-type granules (27.3% increase in *Ttparc6-1 aabb* and 43.5% increase in *Ttparc6 BC aabb*) (Fig. 3d). However, we detected a large, significant increase in B-type granule content in both *Ttparc6-1 aabb* and *Ttparc6 BC aabb* double mutants, that ranged from 67-73% in double mutant genotypes vs. 35-45% in WT and their wild-type segregants (Fig. 3e). There were no significant differences in the numbers of starch granules present (per mg of starch) between the *Ttparc6* double mutants and the controls (Fig. 3f).

**Figure 3:**
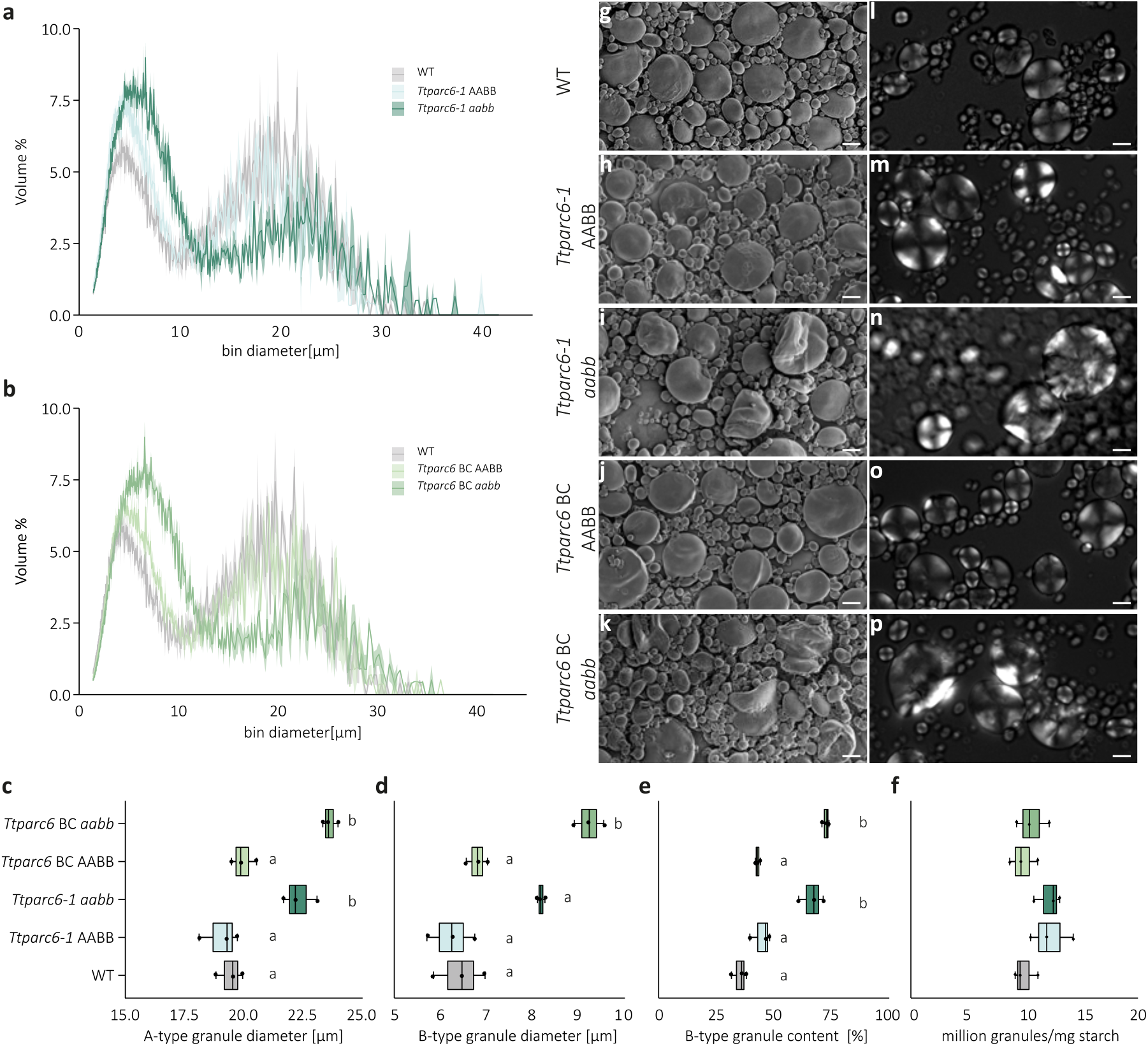
Size distribution and morphology of purified starch granules from mature grains of *Ttparc6* mutants. **(a-b)** Size distribution plots from Coulter counter analysis. The volume of granules at each diameter relative to the total granule volume was quantified using a Coulter counter. Values represent mean (solid line) ± SEM (shading) of three replicates using grains harvested from separate plants. **(c-e)** Granule size parameters obtained from fitting a log-normal distribution to the B-type granule peak and a normal distribution to the A-type granule peak in the granule size distribution data presented in (a – b). Three biological replicates were analysed: **(c)** A-type granule diameter (in μm). Significant differences under a one-way ANOVA and all pairwise multiple comparison procedures (Tukey’s test) are indicated with different letters (p ≤ 0.05). **(d)** B-type granule diameter (in μm). Significant differences under a Kruskal-Wallis one-way ANOVA on the ranks are indicated with different letters (p ≤ 0.019). **(e)** B-type granule content by percentage volume. Significant differences under a one-way ANOVA and all pairwise multiple comparison procedures (Tukey’s test) are indicated with different letters (p ≤ 0.019). **(f)** Granule number per milligram (mg) starch quantified on the Coulter counter. There was no significant difference between genotypes under a one-way ANOVA. **(g-k)** Scanning Electron Microscopy of starch granules from mature grain. Bars = 10 μm. **(l-p)** Polarised light microscopy of starch granules from mature grain. Bars = 10 μm

Interestingly, we also observed a striking dosage effect of *Ttparc6* mutations on the granule size distribution. The *Ttparc6* single homeolog mutants had an intermediate change in granule size distribution, and the associated size parameters (A-type granule diameter, B-type granule diameter, and B-type granule content) were in between those of the double mutants and the wild-type segregants (Fig. S5a-h).

We then examined starch granule morphology of the purified starches using the Scanning Electron Microscope (SEM). Consistent with the results from the size quantification using the Coulter Counter, we observed extremely large A-type granules. Surprisingly, most of the A-type granules in the double mutants had a distinct lobate, crumpled appearance (Fig. 3g-k). Using polarised light microscopy, most of the very large A-type granules had a disrupted Maltese cross, and some had no cross (Fig. 3l-p). A similar lobed surface structure and altered birefringence was observed in A-type granules of the single homeolog mutants (Fig S5i-r).

### Granule morphology is altered throughout grain development in Ttparc6 mutants

To determine how the alterations in granule size and morphology arise in the *Ttparc6* mutants, we examined granules of the *Ttparc6-2* double mutant in developing grains at 12, 16 and 21 days after flowering (DAF) (Figs. 4, 5). At 12 DAF, before the initiation of B-type starch granules, the A-type granules of the *Ttparc6-2 aabb* mutant were similar in size to those of the wild-type (mean diameter of 14.4 µm and 14.0 µm respectively) (Figs. 4a, 5a). However, even at this time point, the double mutant already had strong alterations in A-type granule morphology that were similar to those observed in the mature grain (Figs. 4e, f, m, n). The synthesis of B-type granules had initiated by 16 DAF. By this timepoint, the A-type granules were significantly larger in diameter than the wild-type (5% increase) and B-type granule content was also larger than that of the wild-type (98% increase) (Figs. 4b, 5c). A-type granules in the double mutant remained distinctly lobate, while B-type granules were round and similar to the wild-type (Fig. 4g, h, o, p). The differences in A-type granule diameter, B-type granule diameter and B-type granule content in the *Ttparc6-2 aabb* mutant compared to the wild-type controls increased as grain development progressed (Figs. 4c, d, 5). The mature grains of the double mutant had a similar granule size distribution to those observed in the experiments of Fig. 4 (Figs. 4d, k, l, s, t, 5).

**Figure 4:**
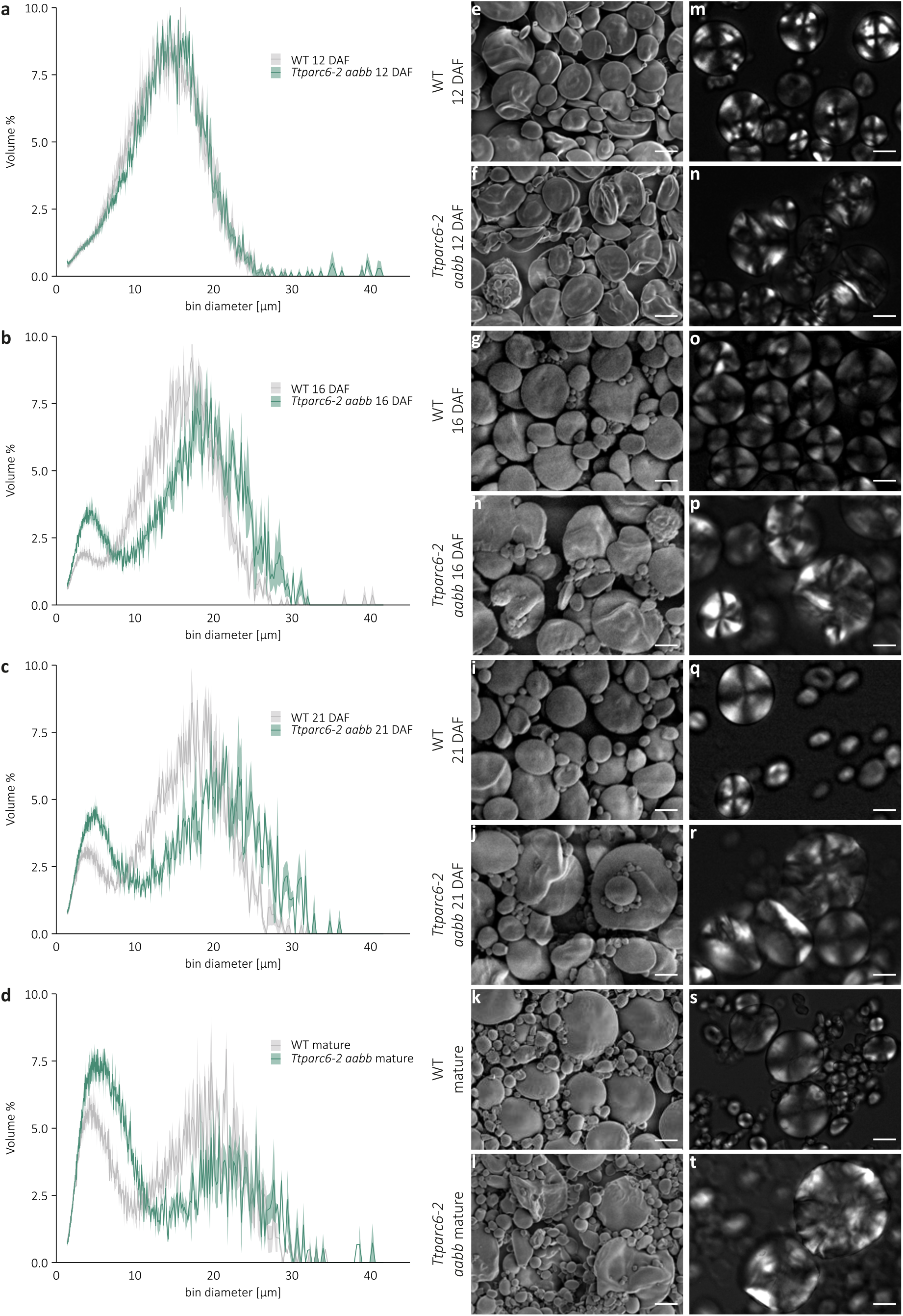
Size distribution and morphology of purified starch granules from developing grains of *Ttparc6-2* mutants. **(a-d)** Size distribution of purified starch granules from developing (12, 16, 21 DAF) and mature grain. The volume of granules at each diameter relative to the total granule volume was quantified using a Coulter counter. Values represent mean (solid line) ± SEM (shading) of three biological replicates using grains harvested from separate plants. **(e-l)** Scanning Electron Microscopy of purified starch granules. Bars = 10 μm. **(m-t)** Polarised light images of purified starch granules. Bars = 10 μm

**Figure 5:**
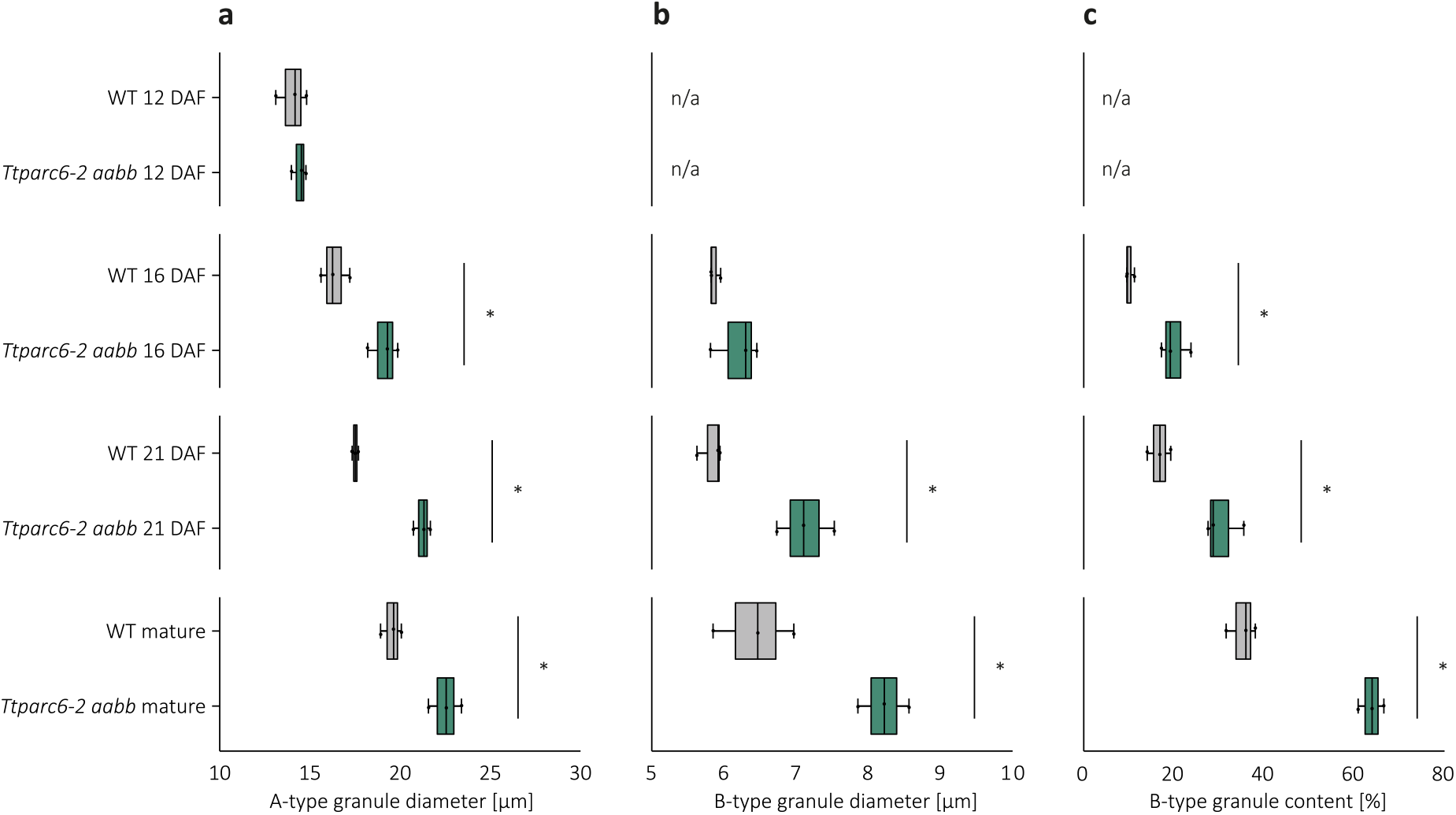
Starch granule size parameters and B-type granule content of developing *Ttparc6-2* grains. **(a-c)** Granule size parameters obtained from fitting a log-normal distribution to the B-type granule peak and a normal distribution to the A-type granule peak in the granule size distribution data presented in (Figure 4). Only A-type granule peaks could be fitted to the distributions at 12 DAF. Three biological replicates from grains harvested from separate plants were analysed, and significant differences (p < 0.05) under a pairwise t-test between genotypes at each timepoint are represented by an asterisk. **(a)** A-type granule diameter (in μm). **(b)** B-type granule diameter (in μm). **(c)** B-type granule content by percentage volume.

### Ttparc6 double mutants have enlarged amyloplasts that contain multiple starch granules

To assess the effects of the *Ttparc6* mutation on endosperm amyloplasts, we examined sections of developing WT and *Ttparc6-2 aabb* grain at 16 DAF using Transmission Electron Microscopy (TEM). In the wild-type, the amyloplast envelope was tightly associated with the large A-type granules (Fig. 6a, b). We did not see protrusions containing additional granules (A- or B-type granules). However, we observed examples of multiple small B-type granules enclosed within a single amyloplast envelope, in vesicle-like compartments (Fig. 6c). We found amyloplast size in the *Ttparc6-2 aabb* mutant was increased compared to the WT. There were single amyloplast compartments containing multiple A- and B-type granules (Fig. 6e). Even in amyloplasts where only a single A-type granule could be observed, the amyloplast envelope was less closely associated with the starch granules than in the wild-type (Fig. 6d). As observed in the wild-type, some amyloplasts in the mutant appeared to contain multiple B-type granules (Fig. 6f).

**Figure 6:**
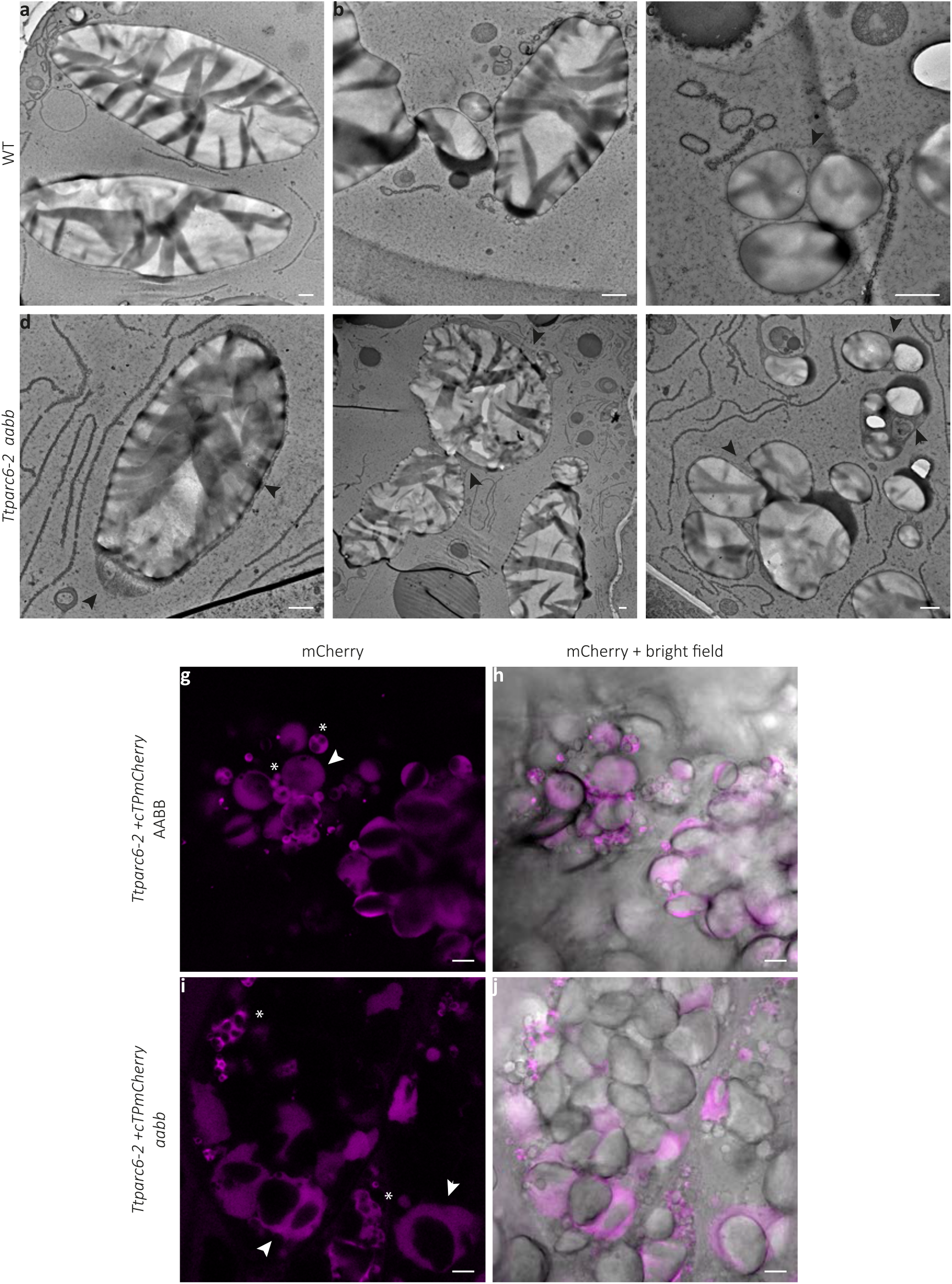
**Amyloplast structure of developing grains (16 DAF) of the *Ttparc6* mutant.** (**a-f)** TEM images of endosperm sections of developing grain at 16 DAF. Arrows indicate amyloplast membrane. Bars = 1 μm. **(g-j)** Confocal laser-scanning imaging of endosperm sections of developing grain at 16 DAF, in lines stably overexpressing the amyloplast marker *cTPmCherry* shown in magenta. Arrows indicate amyloplasts containing A-type starch granules, whereas asterisks point to clusters of B-type starch granules. Bars = 1 μm.

In addition, we used confocal microscopy to examine amyloplasts in segregants of the *Ttparc6-2 aabb* double mutant carrying a fluorescent amyloplast reporter transgene (see above). We imaged cross sections of the *Ttparc6-2 + cTPmCherry aabb* double mutants as well as the *Ttparc6-2 + cTPmCherry* AABB wild-type segregant at 16 DAF. Amyloplast size was drastically increased in the double mutant compared with the wild-type segregant, and many amyloplasts in the mutant contained more than one large A-type granule (Fig. 6). Within these amyloplasts the A-type granules appeared to be separated by stromal space (Fig. 6i, j). Amyloplasts in the wild-type segregant were not observed to contain more than one A-type granule (Fig. 6g, h). Consistent with the TEM images however, there were multiple B-type granules within one amyloplast compartment in both wild-type and mutant (Fig. 6g-j). We verified that the overexpression of cTPmCherry did not influence the *Ttparc6 aabb* phenotype, by confirming that plant growth, grain size, starch content, and granule size distribution were comparable between the lines with and without the reporter (Fig. S6).

### Composition of starch of the Ttparc6 double mutants is similar to wild-type durum wheat starch

We tested whether the highly altered starch granule morphology of the *Ttparc6* mutants resulted from changes in the starch polymer structure and composition. Although increased amylose content relative to the wild type was observed in both the backcrossed and non-backcrossed double mutants and some of the single mutants (Fig. S7a; S3q), the differences in the double mutants were not significant when compared to the wild-type segregant controls and are therefore unlikely to result from *Ttparc6* mutations. The chain length distribution of amylopectin in the *Ttparc6* double mutants was indistinguishable from that of the wild-type and the corresponding wild-type segregants (Fig. S7b). Further, Rapid Visco Analysis (RVA) revealed no consistent difference in viscosity or gelatinisation properties - suggesting that the crystalline structure of the starch granules was unlikely to be altered in the mutant, since changes in crystallinity are usually associated with altered gelatinisation temperature (Fig. S7c-d). In conclusion, the altered starch granule morphology is unlikely to arise from differences in starch composition or polymer structure.

### TaPARC6 interacts with both PDV1 and PDV2 paralogs

In contrast to the *Ttparc6* mutant, the *Ttarc6* double mutant did not show increased chloroplast size in the leaves (Fig. S2). The endosperm starch granule size distribution of the *Ttarc6* double mutant (*Ttarc6 aabb*) was also similar to the wild-type segregant (*Ttarc6* AABB) (Fig. S8a). This raised the possibility that the mechanism of plastid division is different in wheat from that in Arabidopsis, in that *arc6* mutations do not have a strong effect. Alternatively, the position of premature-stop mutations in our wheat *Ttarc6* mutants (in both A and B homeologs) (Fig. S2) might allow the production of a truncated protein with residual function. However, alignment of the amino acid sequences of *Tt*ARC6 and the Arabidopsis *At*ARC6, showed that any putative truncated protein in the *Ttarc6* lines would be terminated at a position that is similar to a previously characterised truncated AtARC6 protein (Atarc6ΔIMS), missing the C-terminal inter-membrane space region (Glynn et al., 2008) (Fig. S9a). In Arabidopsis, this truncation greatly impairs ARC6 function as it removes the C-terminal interaction site of AtARC6 with AtPDV2 and consequently disrupts plastid division (Glynn et al., 2008). We therefore analysed whether the wheat ARC6 and PARC6 proteins are capable of interacting with the wheat PDV2 and PDV1 orthologs respectively, as previously observed for the homologous proteins in Arabidopsis. We identified the corresponding wheat orthologs of PDV1 and PDV2 using BLAST and phylogenetic tree analysis (Fig. S9b). In vascular plants, an early duplication gave rise to both PDV1 and PDV2, proteins. Interestingly, within the Pooidae, there was an additional duplication of PDV1 (resulting in PDV1-1 and PDV1-2) also hinting possible differences in the plastid division mechanism in wheat compared to Arabidopsis (Fig. S9b).

We cloned the A-genome homeologs of wheat *ARC6*, *PARC6, PDV1-1*, *PDV1-2* and *PDV2. Ta*PARC6 and *Ta*ARC6 were N-terminally tagged with a yellow fluorescent protein (YFP) and transiently expressed in *Nicotiana benthamiana* under control of an Arabidopsis Ubiquitin10 promoter and a Cauliflower Mosaic Virus (CaMV) 35S promoter, respectively. Using confocal microscopy, we observed that *Ta*PARC6-A1-YFP localised to distinct puncta in the plastid in pavement cells (Fig. 7a-c). These puncta localised more towards the periphery of the chloroplast and did not colocalise with the chlorophyll autofluorescence, indicating they might be at the chloroplast envelope (Fig. 7a-c). *Ta*ARC6-A1-YFP was also observed at the chloroplast periphery, but did not form puncta (Fig. 7d-f). *Ta*PDV1-1-A1, *Ta*PDV1-2-A1 and *Ta*PDV2-A1 are outer envelope proteins and are targeted towards the chloroplasts and anchored in the outer membrane by a C-terminal sequence rather than an N-terminal transit peptide. Thus, we tagged *Ta*PDV1-1-A1, *Ta*PDV1-2-A1 and *Ta*PDV2-A1 with an N-terminal green fluorescent protein (GFP). We were unable to localise GFP-*Ta*PDV1-1-A1 when transiently expressed in in *Nicotiana benthamiana* under control of a CaMV 35S promoter due to low signal intensity. However, GFP-*Ta*PDV1-2-A1 clearly localised to the chloroplasts, although fluorescence intensity was weak (Fig. 7g-i). GFP-*Ta*PDV2-A1 localised around the chlorophyll fluorescence, indicating a possible localisation to the chloroplast envelope.

**Figure 7:**
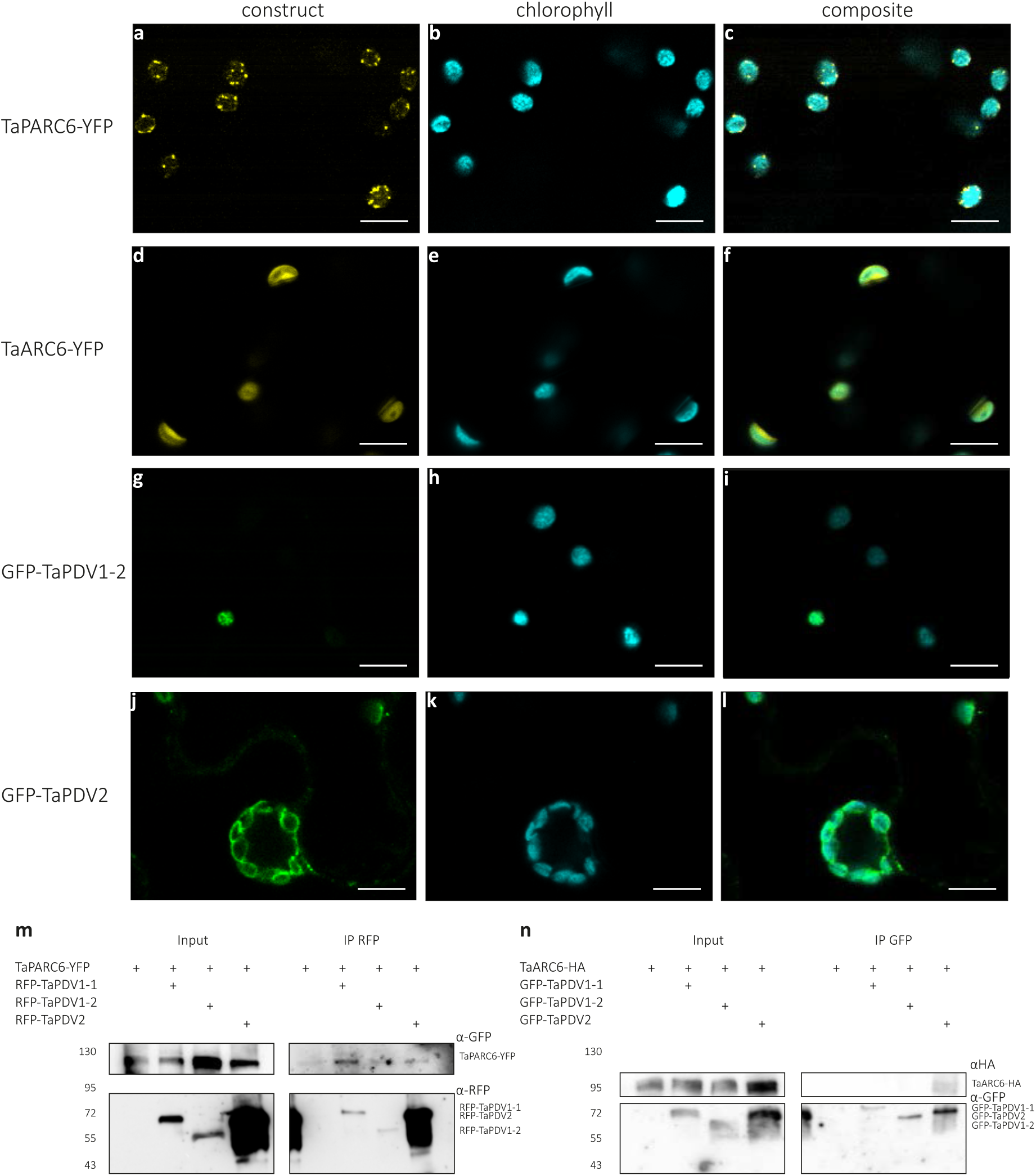
Localisation and co-immunoprecipitation assays of *Ta*PARC6, *Ta*ARC6 and *Ta*PDV isoforms. **(a-l)** Images of transiently expressed *ZmUbi:TaPARC6-YFP*, *CaMV35S:TaARC6-YFP*, *CaMV35S:GFP-TaPDV1-2* and *CaMV35S:GFP-TaPDV2* in *Nicotiana benthamiana* epidermal cells. Images were acquired using confocal laser-scanning microscopy. The YFP and GFP fluorescence are shown in yellow and green, while chlorophyll autofluorescence is shown in cyan. Bars = 10 μm. **(m)** Immunoprecipitation (IP) assay using anti-RFP beads for interactions between TaPARC6-YFP and RFP-TaPDV1-1, RFP-TaPDV1-2 and RFP-TaPDV2, transiently co-expressed in *Nicotiana* leaves. Immunoblots RFP and GFP antibodies were used to detect the proteins. **(n)** Immunoprecipitation (IP) assay using anti-GFP beads for TaARC6-HA and GFP-TaPDV1-1, GFP-TaPDV1-2 and GFP-TaPDV2, transiently co-expressed in *Nicotiana* leaves. Immunoblots HA and GFP antibodies were used to detect the proteins.

In co-immunoprecipitation experiments, *Ta*PARC6-GFP interacted with RFP-*Ta*PDV1-1, and also weakly with RFP-*Ta*PDV1-2 and RFP-*Ta*PDV2 (Fig. 7m). However, *Ta*ARC6-HA only interacted with GFP-*Ta*PDV2 (Fig. 7n), as was previously shown for Arabidopsis (Wang et al., 2017). Therefore, it is possible that in wheat, *Ta*PARC6 might be able to compensate for the loss of *Ta*ARC6 function, by interacting with PDV2 in addition to PDV1-1 and PDV1-2.

## Discussion

### Mutation of TtPARC6 increases amyloplast size and alters starch granule morphology in durum wheat endosperm

Here, we demonstrated that amyloplast architecture, in addition to intrinsic starch polymer properties and granule initiation patterns, is an important factor that determines starch granule morphology. There are numerous examples of altered granule morphology in wheat arising as a consequence of mutations in genes that affect starch polymer biosynthesis and structure (e.g. SS3 and SBE2) or granule initiation patterns (SS4, BGC1, and MRC) (Carciofi et al., 2012; Chia et al., 2020; Hawkins et al., 2021; Chen et al., 2022a; Fahy et al., 2022). However, we achieved highly modified granule morphology after mutating a component of plastid division. Our durum wheat mutants defective in *TtPARC6* not only had increased chloroplast size in leaves but also increased amyloplast size in developing endosperm (Figs. 1d-h, 6). This was accompanied by increased size of both A- and B-type granules in amyloplasts. It is possible that the increases in amyloplast size and accessible stromal volume in the mutant relative to the wild type may facilitate the formation of larger starch granules. Increased granule size in the mutant relative to the wild-type was noticeable at 16 DAF, shortly after the initiation of B-type granules, while at 12 DAF, starch granule size in the *Ttparc6* double mutant was still similar to the wild-type (Figs. 4-5). It is plausible that in early endosperm development, granule size in the wild type is not yet limited by the available space in the amyloplast while at later stages, amyloplast size potentially becomes a limiting factor.

In addition to the increased starch granule size, we observed that the A-type granules of the *Ttparc6* double mutants had drastically altered, lobate granule morphology, compared to the smooth-surfaced disc shape in the wild-type (Fig. 3). This altered morphology manifested early during grain development (12 DAF), even when the granules of the mutant had the same diameter as those of the wild type (Fig. 4). The morphogenesis of wild-type A-type granules during endosperm development was studied in detail by Evers (1971), who reported that A-type granules are initially round, then a grooved annular concretion surrounds two thirds of the granule in an equatorial plane, eventually surrounding the spherical granule as a flange-like outgrowth to form the disc shaped A-type granule (Evers, 1971). It is possible that in the *Ttparc6* mutants, this organised morphogenesis of A-type granule formation is at least partially disrupted. Since amylose content and amylopectin structure were not altered in the *Ttparc6* mutants however (Fig. S7a, b and Fig S3q), the aberrant size and shape of these granules cannot be caused by differences in starch polymer properties. The granules of the mutant also had similar gelatinisation properties (Fig. S7c, d), indicating that crystalline structure is not likely to be altered. Therefore, the disrupted maltese crosses on the A-type starch granules in polarised light are likely caused by increased refraction on the lobate granule surface rather than changes in starch granule crystallinity (Figs. 3k-o, 4m-t). It seems plausible that the enlarged stromal volumes in the *Ttparc6* mutant amyloplasts not only accommodate increased starch granule size, but also influence the usually organised formation of proper A-type granule shape (Fig. 8). Whether this might be due to altered spatial patterns of starch granule growth during granule formation in enlarged plastid compartments remains to be investigated. We recently demonstrated that disrupting the morphology of stromal pockets between the thylakoid membranes in which starch granules form leads to altered granule size and surface structure (Esch et al., 2022), and similar changes in the stromal compartments in *Ttparc6* amyloplasts may explain the altered morphology of the A-type granules (Fig. 3g-p).

**Figure 8:**
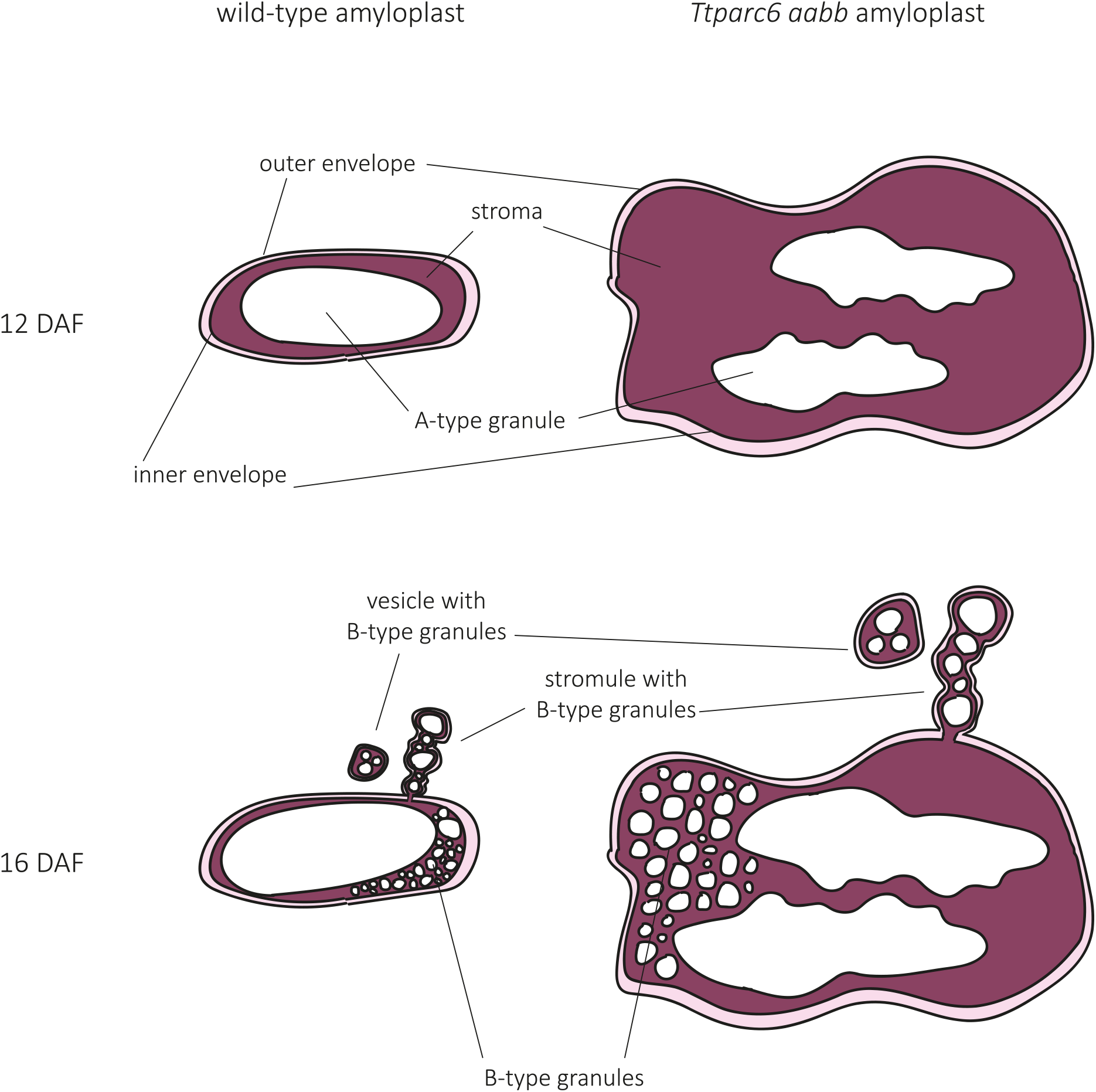
Model of amyloplast and starch granule structures in wild-type and the *Ttparc6* double mutant during endosperm development.

Correct amyloplast size appears to also be important for establishing the proper ratio of A and B-type granule numbers. We observed that increased stromal volume in amyloplasts is associated with greater numbers of starch granules in each amyloplast (Fig. 8). In the mutant, we identified many examples of amyloplasts containing multiple A-type granules, which were not observed in the wild-type (Fig. 6). Also, although the size of both A- and B-type granules was increased in the mutant, the total number of starch granules per milligram purified starch was the same as the wild type (Fig. 3f), which can only be explained by a large relative increase in the number of the smaller B-type granules. This increase in relative number, together with the larger size of individual B-type granules, likely contributed to the higher B-type granule content (as % volume) in *Ttparc6* mutants compared to the wild-type controls (Fig. 3e).

Previously, different models have been proposed regarding the compartmentalisation of B-type granules. It was suggested that B-type granules were initiated and contained in amyloplast stromules (Parker, 1985; Langeveld et al., 2000), or in separate vesicle-like structures (Buttrose, 1960). Our analysis of amyloplast ultrastructure in addition to live-cell imaging of amyloplasts revealed B-type granules in both amyloplast stromules and in separate vesicle-like amyloplasts at 16DAF, in both wild-type and *Ttparc6* double mutants (Fig. 6, 8). In addition, we saw B-type granules within the main compartment that contained the A-type granules in both genotypes (Fig. 6, 8). The occurrence of these different features in wheat supports the hypothesis that stromule formation is an intermediate state of amyloplasts containing B-type granules budding from existing amyloplasts that contain A-type granules, which was recently proposed from similar observations in barley (Matsushima and Hisano, 2019). This is also consistent with early observations that in wheat endosperm, plastid protrusions tended to be short lived (Bechtel and Wilson, 2003).

Our results therefore advance the current models of starch granule formation in wheat by demonstrating that number and size of both A- and B-type starch granules is dependent on amyloplast size and accessible stromal volume (Fig. 8). While this is most pertinent to the bimodal type of starch granules that is unique to the Triticeae, it is also consistent with observations in other systems. In Arabidopsis plastid division mutants, the number of starch granules in the enlarged chloroplasts increases corresponding to the increase in stromal volume such that the number of granules per stromal volume remains similar to the wild-type, and granule size is unaffected (Crumpton-Taylor et al., 2012) (Esch et al., 2022). Moreover, in rice endosperm, various abnormal amyloplast and compound granule morphologies were observed in lines with mutations or silencing in plastid division genes (FtsZ1, FtsZ2-1, PDV1, MinD, MinE, ARC5), including large, elongated, fused, or pleiomorphic amyloplasts (Yun and Kawagoe, 2009; Yun and Kawagoe, 2010). Further, mutations in SSG4 and SSG6, two proteins suggested to be involved in regulation of amyloplast size and development, result in larger amyloplasts in the endosperm (Matsushima et al., 2014; Matsushima et al., 2016; Cai et al., 2022). In all cases examined, the size of the compound granule and the number of the individual granulae per amyloplast was increased. However the individual starch granulae were smaller, and the granulae were irregularly shaped (Yun and Kawagoe, 2009; Yun and Kawagoe, 2010; Matsushima et al., 2014; Matsushima et al., 2016). Taken together with our work, the number of granules per plastid appears to increase with stromal volume, albeit to varying degrees depending on species.

Another common feature of the examined species is that they retain their native granule initiation types (i.e., simple vs. compound) regardless of changes in plastid size. Amyloplast division mutants of rice always produced compound granules; and despite there being multiple A- and B- type starch granules per amyloplast in the wheat *Ttparc6* double mutants, these granules did not fuse or form compound-type starch granules. The formation of compound or bimodal-type starch granules is thus independent from amyloplast size and starch granule number. It was proposed that the formation of compound-type starch granules in rice is potentially dependent on amyloplast sub-compartmentalisation (Yun and Kawagoe, 2010). While its nature is not fully understood, the presence of such compartmentalisation may be more important than amyloplast size for determining different granule types.

### PARC6 in wheat may complement ARC6 deficiency

In Arabidopsis, lack of PARC6 causes diverse plastid morphology phenotypes among different epidermal cell types, which are different and more complex than the phenotype observed in mesophyll cells, where the plastids are consistently increased in size and fewer in number (Ishikawa et al., 2020). Since *Ttparc6* mutants had increased plastid size in both leaves and endosperm, PARC6 appears to be a common element in plastid division in the organs in wheat. However, in strong contrast to the *Ttparc6* double mutants, the *Ttarc6* double mutant had no discernible changes in the size of leaf mesophyll chloroplasts, and also had normal starch granule size distribution in the endosperm (Figs. S2b-c, S8a). This was surprising since in Arabidopsis, the lack of ARC6 causes very strong increases in plastid size, both in leaves and root columella cells (Robertson et al., 1995; Glynn et al., 2008). The mutations in the *Ttarc6* double mutants led to premature stop codons in the coding sequence of both A and B-homeologs, just after the transmembrane domain (Figs. S2a, S9a). If these mutations do not fully knockout protein production, they would at least delete the C-terminal region necessary for interaction with PDV2 (Wang et al., 2017). In Arabidopsis, the deletion of this C-terminal region (like in AtARC6ΔIMS; Fig. S9a) greatly reduced ARC6 function, and could only partially rescue the plastid division phenotype of *arc6* (Glynn et al., 2008). Localisation and co-immunoprecipitation experiments in *Nicotiana benthamiana* indicated that *Ta*ARC6 localises to the chloroplast envelope and can interact with *Ta*PDV2, but not with either of the PDV1 paralogs in wheat (*Ta*PDV1-1 and *Ta*PDV1-2) (Fig. 7). In contrast, *Ta*PARC6 could interact with *Ta*PDV1-1, as well as weakly with *Ta*PDV1-2 and *Ta*PDV2. It is possible that in wheat, the ability of PARC6 to interact with *Ta*PDV2 as well as *Ta*PDV1-1 and *Ta*PDV1-2 allows it to compensate for a loss of ARC6 function, which could explain the lack of plastid-division phenotype in *Ttarc6* mutants.

Despite the potential overlap in interactions, it is likely that *Ta*PARC6 and *Ta*ARC6 retain distinct functions, as reported in Arabidopsis (Zhang et al., 2009; Zhang et al., 2016; Sun et al., 2023). The two proteins showed different subcellular localisations: *Ta*PARC6 formed distinct puncta at the chloroplast envelope, similar to those reported previously for the Arabidopsis ortholog (Glynn et al., 2009; Ishikawa et al., 2020) (Fig. 7a-c). *Ta*ARC6, in contrast, appeared to be homogenously distributed in the chloroplast envelope (Fig. 7d-f). Separate functions of PARC6 and ARC6 in the wheat endosperm is supported by their different temporal patterns of gene expression: *TtPARC6* expression is highest during early endosperm development (6 DPA) and lowest during late developmental stages (20 and 30 DPA) (Fig. S8d-e). For *TtARC6* however, expression in the endosperm is not only about tenfold higher than that of *TtPARC6*, but also peaks at about 15 DPA, which coincides with B-type granule initiation (Fig. S8b-c). Diverse temporal expression patterns were also observed for *Tt*PDV paralogs. Interestingly, while expression of *TtPDV1-2-A1* and *TtPDV1-2-B1* in the developing endosperm mimics the patterns of *TtPARC6*, expression patterns of *TtPDV1-1-A1* and *TtPDV1-1-B1* differed from each other and from *TtPARC6*. *TtPDV1-1-A1* and *TtPDV2-A1* had similar expression patterns to *TtARC6* (Fig. S8f-j). The apparently diverse functions of these PDV paralogs in grasses may be an interesting line of future investigation. Further work is also required to determine if there are mechanistic differences in the function of PARC6 and other plastid division components between the leaves and endosperm, as well as in their regulation. For example, it was recently shown that the interaction between *At*PARC6 and *At*PDV1 is regulated by light, through redox and magnesium (Sun et al., 2023). However, in endosperm amyloplasts of wheat, light is unlikely to be one of the factors promoting interaction of PARC6 and PDV1.

### PARC6 is a novel gene target for modifying wheat starch

Mutation of *PARC6* enabled for the first time, to our knowledge, production of larger starch granules in wheat endosperm. Several benefits of wheat starch with large granule size can be predicted: They include e.g. better milling efficiency, novel functional starch properties and enhanced nutritional properties (Lindeboom et al., 2004; Dhital et al., 2010; Li et al., 2019; Chen et al., 2021). In addition, high B-type granule content is associated with better pasta quality (Soh et al., 2006). Therefore, *PARC6* can be a novel genetic target for modifying starch granule size in wheat. While the feasibility of this will require further field testing, it is promising that under our growth conditions, the *Ttparc6* mutant was not different to the wild type in terms of plant growth and development, photosynthetic efficiency, grain size and yield, and starch content (Figs. 1, 2, S3, S4). This contrasts with the rice *parc6* mutant which had slight reductions in plant growth and grain weight (Kamau et al., 2015). Interestingly, all changes in starch granule morphology in the *Ttparc6* double mutants were less severe in the single mutants, and this dosage effect can be exploited to achieve a range of different starch granule sizes.

## Methods

### Plant material and growth conditions

Mutants in *Triticum turgidum* (cv. Kronos) from the wheat TILLING mutant resource (Krasileva *et al*., 2017) were: Kronos1265 (K1265; C/T at chromosome 2A coordinate 759829502) for *TtPARC6-A1*, Kronos2369 (K2369; G/A at chromosome 2B coordinate 775170338) for *TtPARC6-B1*, Kronos3404 (K3404; C/T at chromosome 6A coordinate 35481841) for *TtARC6-A1* and Kronos2205 (K2205; G/A at chromosome 6B coordinate 64929650) for *TtARC6-B1*. The mutations were genotyped using KASP v4 master mix (LGC) using the primers in Table S1.

Wheat plants were grown in controlled environment rooms or glasshouses. Controlled environments were set to 16 h light/8 h dark cycles with light intensity set to 300-400 µmol photons m^-2^ s^-1^. Glasshouses were set to provide a minimum of 16 h light at 300-400 µmol photons m^-2^ s^-1^. In both cases, temperature was set to 20°C in light and 16°C in dark, and relative humidity was set to 60%. *Nicotiana benthamiana* plants were grown in the glasshouse set to a minimum of 16 h light at 22°C.

### Cloning and construct assembly

To generate the transgenic wheat amyloplast reporter lines, we modified a construct design from Matsushima and Hirano (2019), to use a codon-optimised *mCherry* coding sequence (rather than GFP in the original citation) downstream of the *OsWaxy* transit peptide sequence (sequence in Table S2). This fusion sequence, flanked by attB1 and attB2 recombination sites, was synthesised as a gBlocks fragment (IDT) and recombined into the Gateway entry vector pDONR221 using Gateway BP clonase II (Invitrogen, Thermo Fisher Scientific). The *cTPmCherry* coding sequence was then recombined using Gateway LR clonase II (Invitrogen, Thermo Fisher Scientific) into a modified *pGGG* vector (Hayta et al., 2021), pGGG_AH_Ubi_GW_NosT, encoding for a Hygromycin resistance gene driven by an actin promoter (AH), a gateway cassette for gateway recombination (GW) downstream of the *ZmUbiquitin* promoter (Ubi) and upstream of a Nos terminator (NosT).

*TaPARC6-A1*, *TaARC6-A1*, *TaPDV1-1-A1*, *TaPDV1-2-A1* and *TaPDV2-A1* sequences were obtained from the RefSeq 1.1 genome from Ensembl Plants (Yates et al., 2022). Codon-optimised coding sequences of *TaARC6-A1*, *TaPDV1-1-A1*. *TaPDV1-2-A1* and *TaPDV2-A1*, flanked by attB1 and attB2 recombination sites, were ordered as a gBlocks fragments (IDT) (Table S2). These coding sequences were recombined into pDONR221 as above. A codon-optimised sequence of *TaPARC6-A1,* flanked by attL recombination sites and MluI restriction sites (Table S2), was ordered from Genewiz in a pUC-GW-Kan vector. *TaPARC6-A1* and *TaARC6-A1* were recombined into Gateway expression vectors pUBC-YFP, pB7YWG2 and pJCV52; and *TaPDV1-1_A1*, *TaPDV1-2_A1* and *TaPDV2_A1* were recombined into Gateway destination vectors pK7WGF7 and pGWB555 using Gateway LR clonase II. All constructs were confirmed by Sanger sequencing.

### Plant transformation

The *cTPmCherry* construct was transformed into *T. turgidum* cv. Kronos using Agrobacterium-mediated transformation of embryonic calli, as described in Hayta et al., (2021). Lines with single insertions were selected using RT-PCR against the Hygromycin marker gene (performed by iDNA Genetics, Norwich, UK). *Nicotiana benthamiana* plants were transiently transformed as described in SEP S3.

### Grain and plant morphometrics

The number of grains harvested per plant, as well as grain size traits (area, length, width, total grain weight per plant and thousand grain weight) were quantified using the MARViN seed analyser (Marvitech GmbH, Wittenburg). Grains of 3 plants per genotype (–0 - 259 individual grains per plant) were analysed. The number of tillers were counted in mature plants before grain harvesting.

### Starch purification, granule morphology and size distribution

Starch purification, scanning electron microscopy and polarised light microscopy was performed as described in Hawkins et al. (2021) and SEP S4. Granule size distribution was analysed and plotted in relative volume/diameter using the Multisizer 4e Coulter counter (Beckman Coulter) (SEP S4).

### Total starch content, starch composition, amylopectin structure and Rapid Visco Analysis

Grain starch quantification was performed as described in Hawkins et al. (2021) and SEP S5. Amylopectin structure and amylose content was analysed using purified starch as described in Chen et al. (2022a) and SEP S4. Rapid Visco Analysis (RVA) was carried out on an RVA Tecmaster instrument (Perten) running the pre-installed general pasting method (AACC Method 76-21). Analyses were performed with 1.5 g purified starch or 5 g flour (SEP S4) in 25 mL of water.

### Microscopic analysis of plastid morphology

For the analysis of mesophyll chloroplast morphology: separation of mesophyll cells was performed according to Pyke and Leech (1991) with adjustments described in SEP S6. Mesophyll chloroplasts were imaged using the LSM800 (Zeiss) or the TCS SP8X (Leica) using a 40.0x or 63.0x water immersion objective. Chlorophyll autofluorescence was excited using a white light laser set to 555 nm, 576 nm or 630 nm and emission was detected at 651 nm to 750 nm using a hybrid detector or Airyscan.

For the analysis of endosperm amyloplast morphology using confocal microscopy: Developing grain of amyloplast reporter lines (see above) were harvested at 16 DAF, embedded in 4% low melting agarose and sectioned into 150 µm cross sections using the vibratome VT1000s (Leica). Images were acquired immediately after sectioning on the LSM800 using a 63.0 x oil immersion objective (Zeiss). mCherry signal was excited at 561 nm and emission was detected at 562 nm to 623 nm (605 nm).

For analysis of endosperm amyloplast morphology using transmission electron microscopy (TEM), samples were prepared and imaged as described in Chen et al. (2022a) and SEP S7. All images in this manuscript were processed using the ImageJ software (http://rsbweb.nih.gov/ij/) and Adobe Photoshop 2020. Chloroplast images were additionally processed using the Zeiss ZEN software.

### Protein localisation in Nicotiana benthamiana

*Ta*PARC6-YFP, *Ta*ARC6-YFP, GFP-*Ta*PDV1-2 and GFP-*Ta*PDV2 were localised in *Nicotiana benthamiana* using confocal microscopy as described in SEP S8.

### Protein extraction and Immunoblotting

For the pairwise immunoprecipitation assays, two 1 cm diameter leaf discs from two *Nicotiana benthamiana* leaves transiently expressing the tagged proteins were homogenised in extraction buffer (50mM Tris-HCl, pH 8.0, 150 mM NaCl, 1% v/v Triton X-100, 1x protease inhibitor cocktail, 1 mM DTT). Homogenates were spun at 20,000g, 10 mins and proteins were collected in the supernatant (Input sample). Immunoprecipitation was performed on the input sample using the µMACS GFP Isolation Kit (Miltenyi Biotec) or the RFP-Trap Magnetic Particles (Chromotek) and µMACS Columns (Miltenyi Biotec). For immunoblotting antibodies were used in the following concentrations: 1:5000 anti-GFP (TP401, Torrey pines), 1:2000 anti-RFP (ab34771, Abcam), and 1:5000 anti-HA (ab9110; Abcam). Bands were detected using the anti-rabbit IgG (whole molecule)-Peroxidase (A0545, Sigma) at 1:20000 dilution and the SuperSignal West Femto Trial Kit (Thermo Scientific).

### Gene expression analysis

Normalised values for gene expression (in transcripts per million) in the developing endosperm of *T. turgidum* cv. Kronos were retrieved from Chen et al. (2022b).

## Supporting information

Supplemental Experimental Procedures

## Author contributions

Conceived and designed the experiments: L.E, A.M.S, D.S. Performed the experiments: L.E Q.Y.N, J.E.B, S.H, M.A.S. Analysed data: L.E, Q.Y.N, R.M. Wrote the paper: L.E and D.S (with input from all authors).

## Acknowledgements

The authors thank the John Innes Centre (JIC) Horticultural Services for providing growth facilities and maintenance of plant material, JIC Bioimaging for providing access to microscopes, JIC Crop Transformation for providing transformation resources and expertise, the JIC Genotyping platform for providing DNA extraction and KASP genotyping, and Alexander Watson-Lazowski (Harper Adams University) and Richard Vath (University of Cambridge) for helpful advice on the gas exchange experiments. L.E and D.S are coinventors on a patent for using PARC6 to alter starch granule morphology. We have no other conflicts of interest to declare.

