## Supplemental Experimental Procedures for "Modification of amyloplast size in wheat endosperm through mutation of PARC6 affects starch granule morphology"

**Supplemental Experimental Procedures (SEP)**

*S1: Phylogenetic analysis and gene models*

PARC6, ARC6, PDV1 and PDV2 protein sequences were retrieved from Ensembl Plants and Phytozome (Goodstein et al., 2012; Yates et al., 2022). Proteins were aligned using ClustalW in MEGA7. Phylogenetic analysis was conducted in MEGA7 (Kumar et al., 2016) using the Maximum Likelihood method based on the JTT matrix-based model (Jones et al 1992). Initial tree(s) for the heuristic search were obtained automatically by applying Neighbour-Join and BioNJ algorithms to a matrix of pairwise distances estimated using a JTT model, and then selecting the topology with superior log likelihood value.

Gene models were taken from Ensembl Plants and domains were annotated using Interpro (Yates et al., 2022; Paysan-Lafosse et al., 2023).

*S2: Gas exchange*

Gas exchange measurements were made using an LI-6800P portable photosynthesis system (Li-COR) 40-46 days after germination on the fully expanded flag leaves in the glasshouse ($CO_2$ concentration ca. 412 ppm, light intensity ca. 280 µmol $m^{-2}$ $s^{-1}$, temperature ca 21°C) as described in Watson-Lazowski *et al.* (2022). The responses of the $CO_2$ assimilation rate to step increases in light intensity (AQ) were measured under constant $CO_2$ conditions (412 ppm). AQ measurements were taken after acclimation of 60-120s at increasing light intensities (0, 20, 50, 75, 100, 150, 200, 500, 750, 1000, 1200, 1500, 1800, 2000 µmol $m^{-2}$ $s^{-1}$). The response of the $CO_2$ assimilation rate to step increases of intra-cellular $CO_2$ (A/Ci) was measured at saturating light (2000 µmol $m^{-2}$ $s^{-1}$). The A/Ci curves measured at decreasing and increasing $CO_2$ steps of 400, 300, 200, 100, 50, 0, 400, 400, 600, 800, 1000, 1200 ppm. Maximum rates of carboxylation (Vcmax) and electron transport (Jmax) were calculated from A/Ci curves using the ‘Plantecophys’ package in R by fitting the raw data to a Farquhar, von Caemmerer, and Berry photosynthesis model (Farquhar et al., 1980; Duursma, 2015). AQ and A/Ci curves were measured consecutively. Before carrying out A/Ci curves, leaves were allowed to stabilise for 20 min at maximal light intensity (2000 µmol $m^{-2}$ $s^{-1}$). All measurements were taken 8–14 h after the end of the night.

S3: *Transient transformation of Nicotiana benthamiana*

*Nicotiana benthamiana* plants were transiently transformed using *Agrobacterium tumefaciens* (GV3101) carrying the respective constructs. The bacteria were grown at 28°C for 48 h. Cultures were resuspended in MMA buffer (10 mM MES pH 5.6, 10 mM $MgCl_2$, 0.1 mM acetosyringone) at an optical density of 1.0 at 600 nm for confocal microscopy and of

0.3 (0.2 for p19) at 600 nm for protein extraction, and infiltrated into the abaxial side of the leaf using a syringe. Leaves were harvested for confocal microscopy and protein extraction 48-72 h after infiltration.

*S4: Starch purification, scanning electron microscopy and polarised light microscopy.*
For mature grains, three grains per sample were soaked overnight in double distilled water (dd$H_2O$) at 4°C, then homogenized in a mortar and pestle with additional dd$H_2O$. Developing grains were snap frozen in liquid nitrogen at harvest and stored at -80°C. Seeds were thawed immediately before endosperm dissection, and endosperms were homogenized in dd$H_2O$ using a ball mill at 30 Hz for 1.5 minutes. For large amounts of starch, mature grains were first milled into flour (Cyclone Mill Twister, Retsch). Homogenates were filtered through a 100 µm nylon mesh, centrifuged and the pellet was resuspended in 90% (v/v) Percoll, 50 mM Tris-HCl, pH 8. The suspension was centrifuged at 2500 g for 5 min and the pellet was washed twice in 50mM Tris-HCl, pH 6.8, 10 mM ethylenediaminetetraacetic acid (EDTA), 4% sodium dodecyl sulfate (SDS) (v/v), 10 mM dithiothreitol (DTT). The starch pellet was washed and resuspended in dd$H_2O$.

Granule size distribution was analysed and plotted in relative volume/diameter using the Multisizer 4e Coulter counter (Beckman Coulter). fitted with a 70 µm aperture, operating on either total count mode (measuring a minimum of 500,000 particles) or volumetric mode (measuring a minimum of 1 mL starch suspension). Measurements were conducted with logarithmic bin spacing and were corrected for bin width for presentation on a linear x-axis. A- and B-type granule diameters as well as B-granule contents were extracted by fitting distribution models to the data. Python script available at: https://github.com/DavidSeungLab/Coulter-Counter-Data-Analysis.

The morphology of starch granules was examined by scanning electron microscopy, using a Nova NanoSEM 450 (FEI) scanning electron microscope and the Leica DM6000 microscope for polarised light microscopy. Images were processed using ImageJ software (http://rsbweb.nih.gov/ij/) and Adobe Photoshop 2020.

*S5: Total starch quantification, starch composition and amylopectin structure*
Grain starch quantification was performed using the Total Starch Assay kit (K-TSTA; Megazyme): Flour (milled in ball mill: 5-10 mg) was suspended in 20 µL 80% ethanol and incubated with 500 µL thermostable α-amylase in 100 mM sodium acetate buffer, pH 5, at 99 °C and 1400 rpm for 7 min. Amyloglucosidase was added and incubated at 50 °C and 1000rpm for 35 min. Samples were centrifuged at 20.800g for 10 min and glucose content

was measured in the supernatant using the hexokinase/glucose-6-phosphate dehydrogenase assay (Roche, Basel, Switzerland) to calculate starch content in glucose equivalents.

Amylose content was determined using an iodine-binding method on starch granules dispersed in water, adapted from Washington et al., (2000). Briefly: 1 mg of purified starch (as in S4) was resuspended in 200 µL water, mixed with 200 µL 2 M NaOH solution and incubated at room temperature over night. The starch slurry was neutralised with 400 µL 1 M HCl. 5 µL of the starch suspension were diluted in 220 µL water and 25 µL Lugol solution (Sigma Life Science). Absorbance was measured at 620 nm and 535 nm and Amylose content was calculated as described in Washington et al., (2000). Amylopectin chain length distribution was quantified using High Performance Anion Exchange Chromatography with Pulsed Amperomatric Detection (HPAEC-PAD) on a Dionex ICS-5000-PAD fitted with a PA-100 column (Thermo). The preparation of debranched samples was carried out as described in Streb et al., (2008).

*S6: Cell separation assay*

Leaf segments of the leaf tip of the 3$^{rd}$ fully developed leaf were harvested into 10% formaldehyde solution (Sigma) in PBS (v/v) and incubated in the dark for 2 h. Formaldehyde solution was replaced by 0.1 M $Na_2EDTA$, pH 9 and samples were incubated at 100 rpm and 60 °C for 2 h. Cells were separated by carefully knocking the coverslip during mounting.

*S7: Transmission electron microscopy of developing grain*

Developing grain (16 DAF) were harvested into 2.5% glutaraldehyde in 0.05 M sodium cacodylate, pH 7.4. Samples were post-fixed in 1% (w/v) osmium tetroxide ($OsO_4$) in 0.05 M sodium cacodylate for 2 h at room temperature, dehydrated in ethanol and infiltrated with LR White resin (Agar Scientific, Stansted, UK), using a EM TP embedding machine (Leica, Milton Keynes, UK). LR White blocks were polymerised at 60°C for 16h. For transmission electron microscopy (TEM) ultrathin sections (ca. 90 nm) were cut with a diamond knife and placed onto formvar and carbon coated copper grids (EM Resolutions, Sheffield, UK). The sections were stained using 2% (w/v) uranyl acetate for 1 h and 1% (w/v) lead citrate for 1 min, washed in water and air dried. Sections were imaged on a Talos 200C TEM (FEI) at 200 kV and a OneView 4K x 4K camera (Gatan, Warrendale, PA, USA).

*S8: Protein localisation in Nicotiana benthamiana*

For localisation of flurophore tagged *Ta*PARC6-YFP, *Ta*ARC6-YFP, GFP-*Ta*PDV1-2 and GFP-*Ta*PDV2 in *Nicotiana benthamiana*: Images were acquired on the Leica Stellaris 8 laser scanning confocal microscope using a 40.0x water immersion objective. YFP signal was excited using a white light laser set to 514 nm and emission was detected at 519 nm to 560 nm. GFP signal was excited using a white light laser set to 488 nm and emission was detected at 562 nm to 623 nm. Chlorophyll autofluorescence was excited using a white light laser set to 555 nm, 576 nm or 587nm and emission was detected at 642 nm to 750 nm.
