## Supplemental Figures and Tables for "Modification of amyloplast size in wheat endosperm through mutation of PARC6 affects starch granule morphology"

### Supporting Information

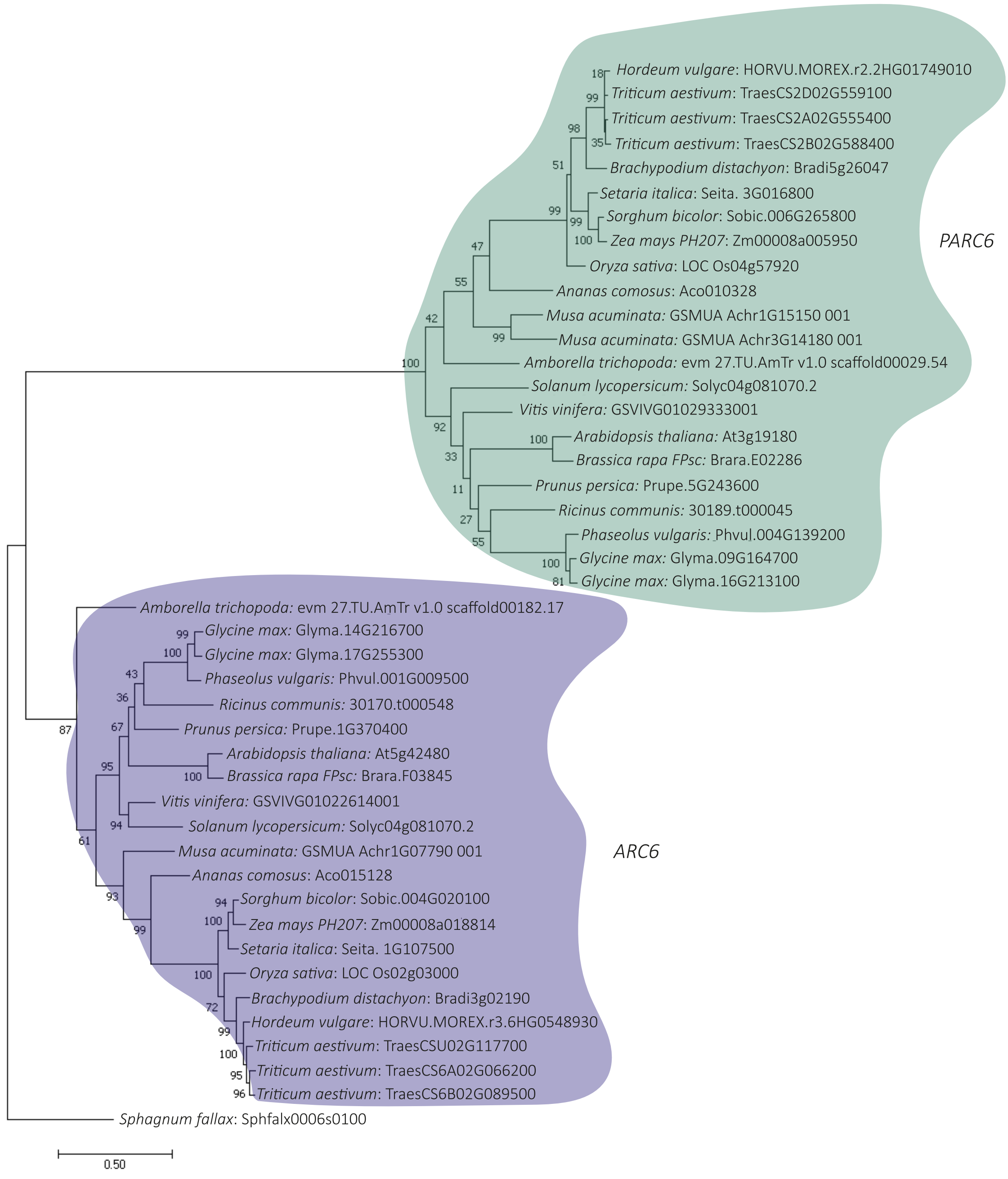


**Figure S1: Molecular phylogenetic analysis of *ARC6* and *PARC6* gene families.**
The tree with the highest log likelihood (-36614.75) is shown. The percentage of trees out of 1000 bootstraps in which the associated taxa clustered together is shown next to the branches. The tree is drawn to scale, with branch lengths and scale bar representing the number of substitutions per site. Full experimental procedures can be found in SEP S1.

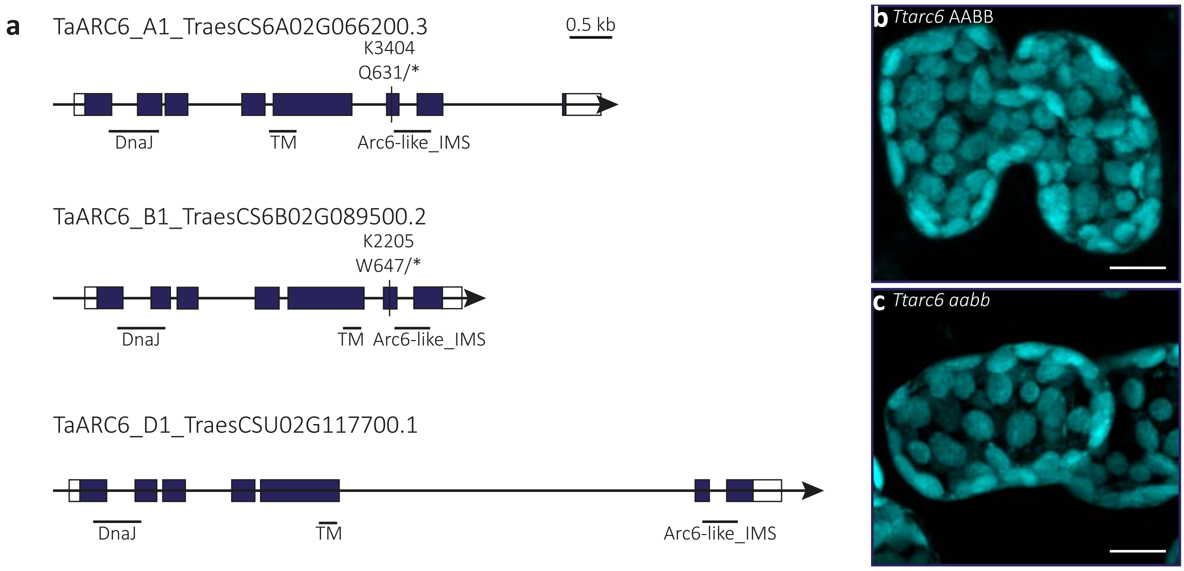


**Figure S2: *TaARC6* gene models and chloroplast morphology of the *Ttarc6* mutant.**
**(a)** Schematic illustration of the gene models for the canonical transcripts of *TaARC6-A1*, *-B1* and *-D1* in bread wheat. Exons are represented as purple boxes and UTRs are represented as white boxes. Mutation sites in K3404 and K2205 are indicated by black lines and the resulting amino acid to stop codon (*) substitutions are annotated. Regions encoding domains are indicated by black horizontal lines (TM: Transmembrane, IMS: Inter Membrane Space).
**(b-c)** Images of mesophyll-cell chloroplasts in the third leaf of *Ttarc6* mutants seedlings. Images were acquired using confocal microscopy and are Z-projections of image stacks. Chlorophyll auto-fluorescence of the chloroplasts is shown in cyan. Bars = 10 μm.

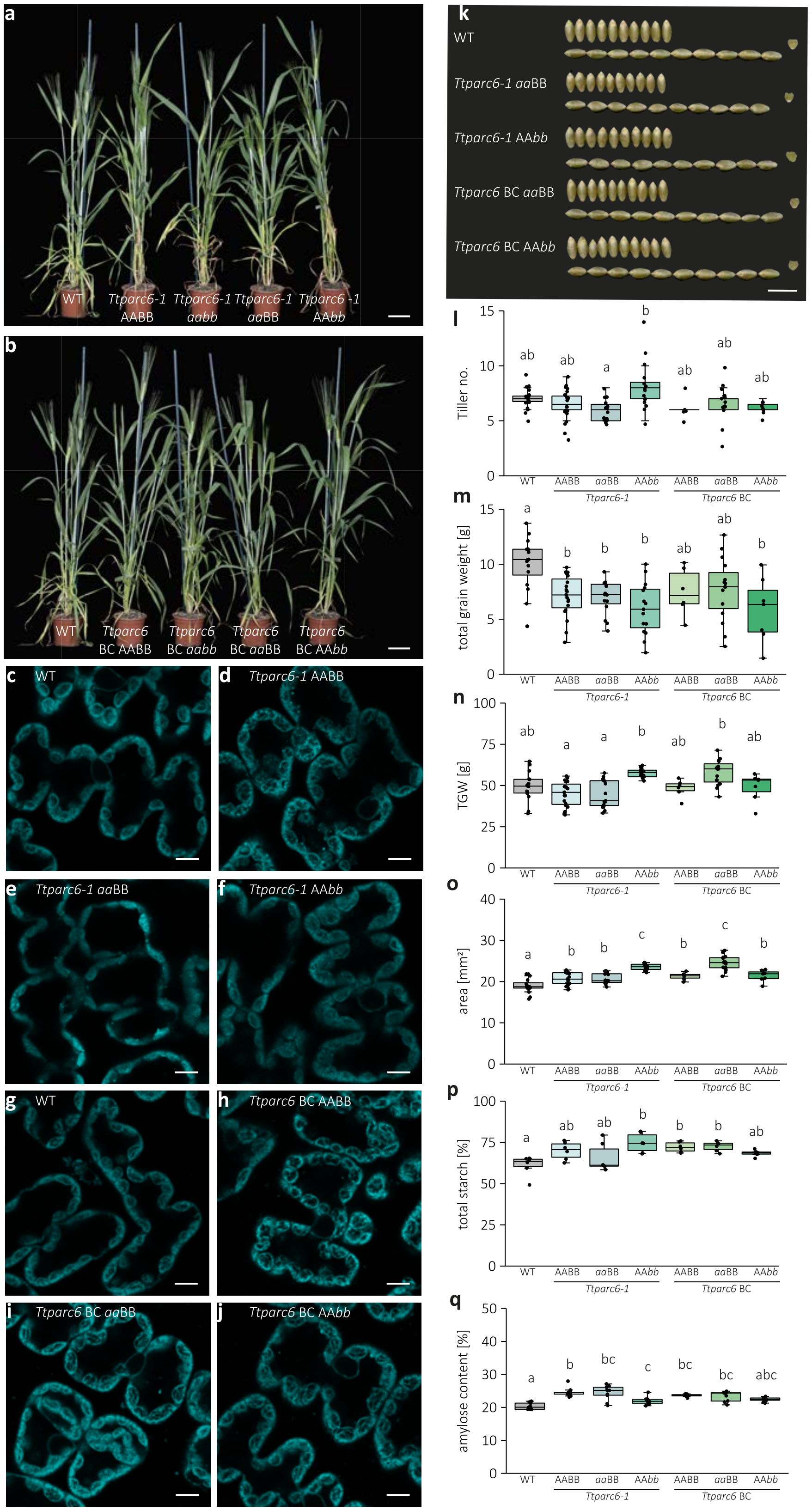


**Figure S3: Growth and seed phenotype of *Ttparc6* single homeolog mutants.**
**(a, b)** Photographs of 8-week-old *Ttparc6* double and single mutants (*Ttparc6-1 aabb*, *Ttparc6-1 aa*BB and *Ttparc6-1* AA*bb*) and the corresponding negative segregant (*Ttparc6-1* AABB); and *Ttparc6* backcrossed double and single mutant (*Ttparc6* BC *aabb*, *Ttparc6* BC *aa*BB and *Ttparc6* BC AA*bb*)

and the corresponding negative segregant (*Ttparc6* BC AABB) and WT wheat (cv Kronos) plants. Bars = 10 cm.
**(c-j)** Images of mesophyll-cell chloroplasts in the third leaf of *Ttarc6* mutant seedlings. Images were acquired using confocal microscopy. Chlorophyll auto-fluorescence in the chloroplasts is shown in cyan. Bars = 10 µm.
**(k)** Photograph of 10 representative mature grains per genotype. Bar = 1 cm.
**(l)** The number of tillers per plant (Tiller no.) of mature plants (n = 6 - 19 per genotype). Significant differences between the lines as determined by Kurskal-Wallis One Way ANOVA on the Ranks all pairwise multiple comparison (Dunn's Method) ($p \leq 0.009$) are represented by different letters.
**(m)** Total grain weight harvested per plant (in g). Dots represent the total grain weight of individual plants (n = 6-19). Significant differences under a one-way ANOVA and all pairwise multiple comparison procedures (Tukey's test) are indicated with different letters ($P \leq 0.001$).
**(n)** Thousand grain weight (TGW) (in g). Dots represent calculated TGW of individual plants (*n* = 7-19) per genotype. Significant differences under a one-way ANOVA and all pairwise multiple comparison procedures (Tukey's test) are represented by different letters ($p \leq 0.001$).
(**o**) Grain size measured as seed area (in $mm^2$). Dots represent measurements for seeds of 3-19 plants per genotype. Significant differences under one-way ANOVA and all pairwise multiple comparison procedures (Tukey's test) are represented by different letters ($p \leq 0.05$).
**(p)** Total starch content as % (w/w). 3 technical replicates of 2 biological replicates per genotype. Significant differences under a one-way ANOVA and all pairwise multiple comparison procedures (Tukey's test) are represented by different letters ($p \leq 0.05$).
**(q)** Amylose content [% of total starch]. Dots represent 3 technical replicates of 3 biological replicates. Significant differences between the lines as determined by a one-way ANOVA on the ranks and all pairwise multiple comparison (Tukey's test) are represented by different letters ($p \leq 0.001$).
For all boxplots, the bottom and top of the box represent the lower and upper quartiles respectively, and the band inside the box represents the median. The ends of the whiskers represent values within 1.5x of the interquartile range, whereas values outside are outliers.

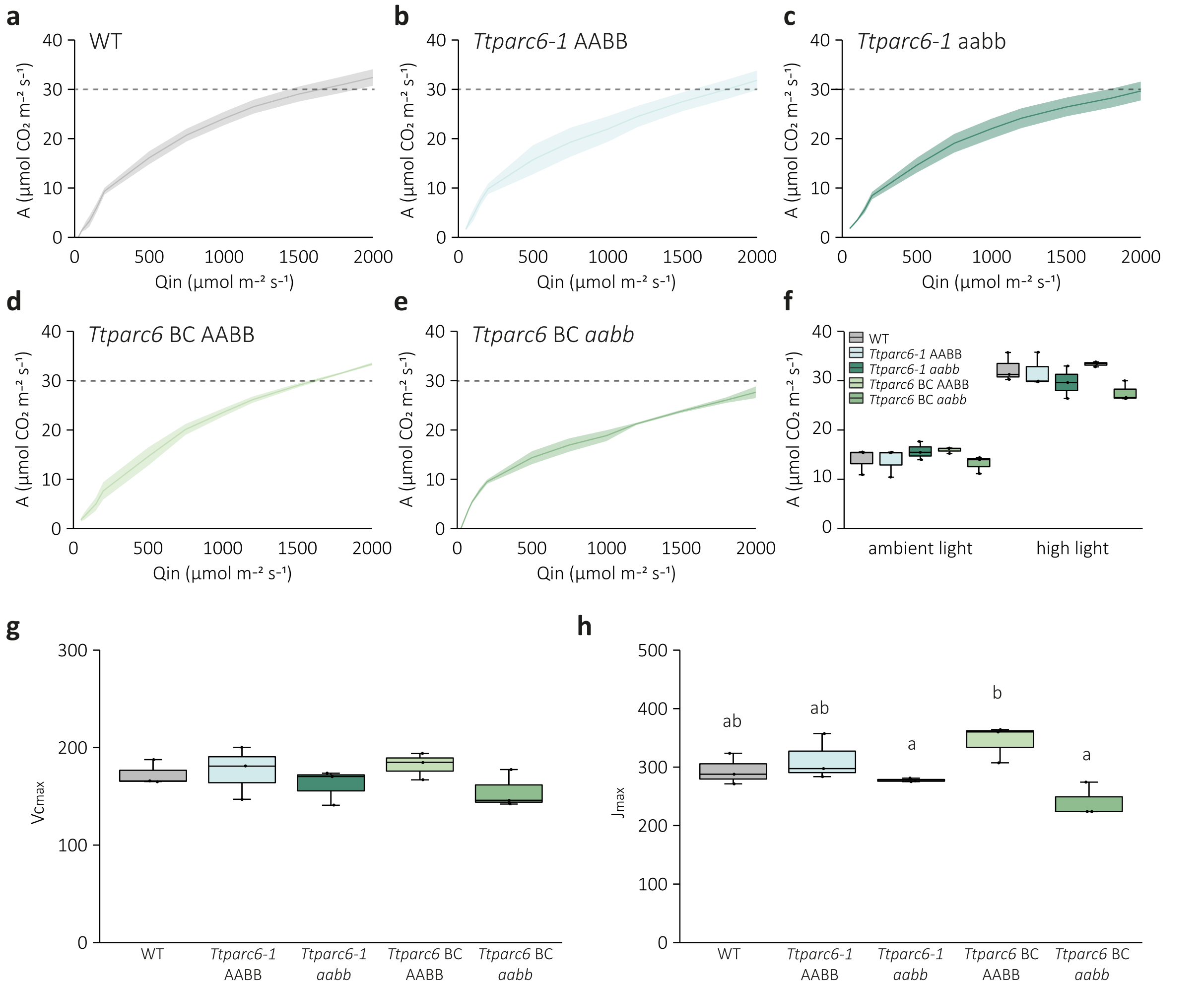


**Figure S4: Photosynthesis parameters of *Ttparc6* mutant plants.**
Light response and A/Ci curves were measured on the flag leaf of 40-46 day old plants in three separate plants (n = 3). Experimental procedures are in SEP S2.
**(a-e)** Light response curves were measured at ambient $CO_2$ levels (412 µmol $m^{-2}$ $s^{-1}$). Lines represent the average and ribbons the standard error. The dotted line at A = 30 µmol $m^{-2}$ $s^{-1}$ is provided to aid comparison between genotypes.
**(f)** Assimilation rate at ambient light (280 µmol $m^{-2}$ $s^{-1}$) and high light (2000 µmol $m^{-2}$ $s^{-1}$).
**(g)** Estimation of Vcmax was extracted from the A/Ci curves using plant ecophys.package (R).
**(h)** Estimation of Jmax was extracted from the A/Ci curves using plant ecophys.package (R).

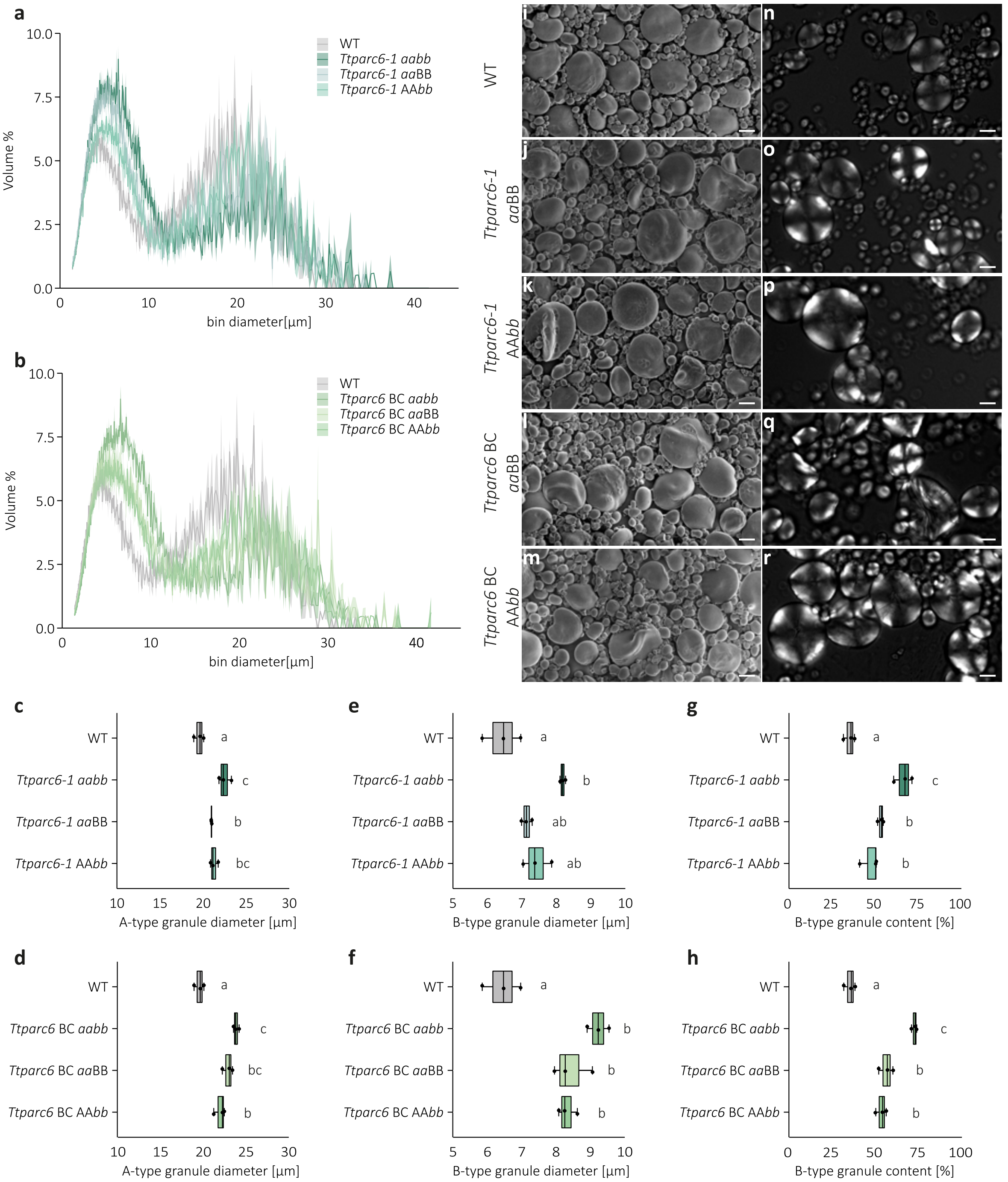


**Figure S5: Size distribution of purified starch granules from mature grains of the *Ttparc6* single mutants.**

**(a-b)** Size distribution plots from Coulter counter analysis. The volume of granules at each diameter relative to the total granule volume was quantified using a Coulter counter. Values represent mean (solid line) ± SEM (shading) of three replicates using grains harvested from separate plants.

**(c-h)** Granule size parameters obtained from fitting a log-normal distribution to the B-type granule peak and a normal distribution to the A-type granule peak in the granule size distribution data presented in (a – b). Three biological replicates were analysed: **(c, d)** A-type granule diameter (in µm). Significant differences under a one-way ANOVA and all pairwise multiple comparison procedures (Tukey's test) are indicated with different letters ($p \leq 0.05$). **(e, f)** B-type granule diameter (in µm). Significant differences under a Kruskal-Wallis one-way ANOVA on ranks and all pairwise multiple comparison procedures (Tukey's test) are indicated with different letters ($p \leq 0.022$) for *Ttparc6-1* lines. Significant differences under one-way ANOVA and all pairwise multiple comparison procedures (Tukey's test) are indicated with different letters ($p \leq 0.05$) for *Ttparc6* BC lines. **(g, h)** B-

type granule content by percentage volume. Significant differences under a one-way ANOVA and all pairwise multiple comparison procedures (Tukey's test) are represented with different letters ($p \leq 0.05$).
**(i-m)** Scanning Electron Microscopy of purified starch granules. Bars = 10 μm.
**(n-r)** Polarised light microscopy of purified starch granules. Bars = 10 μm.

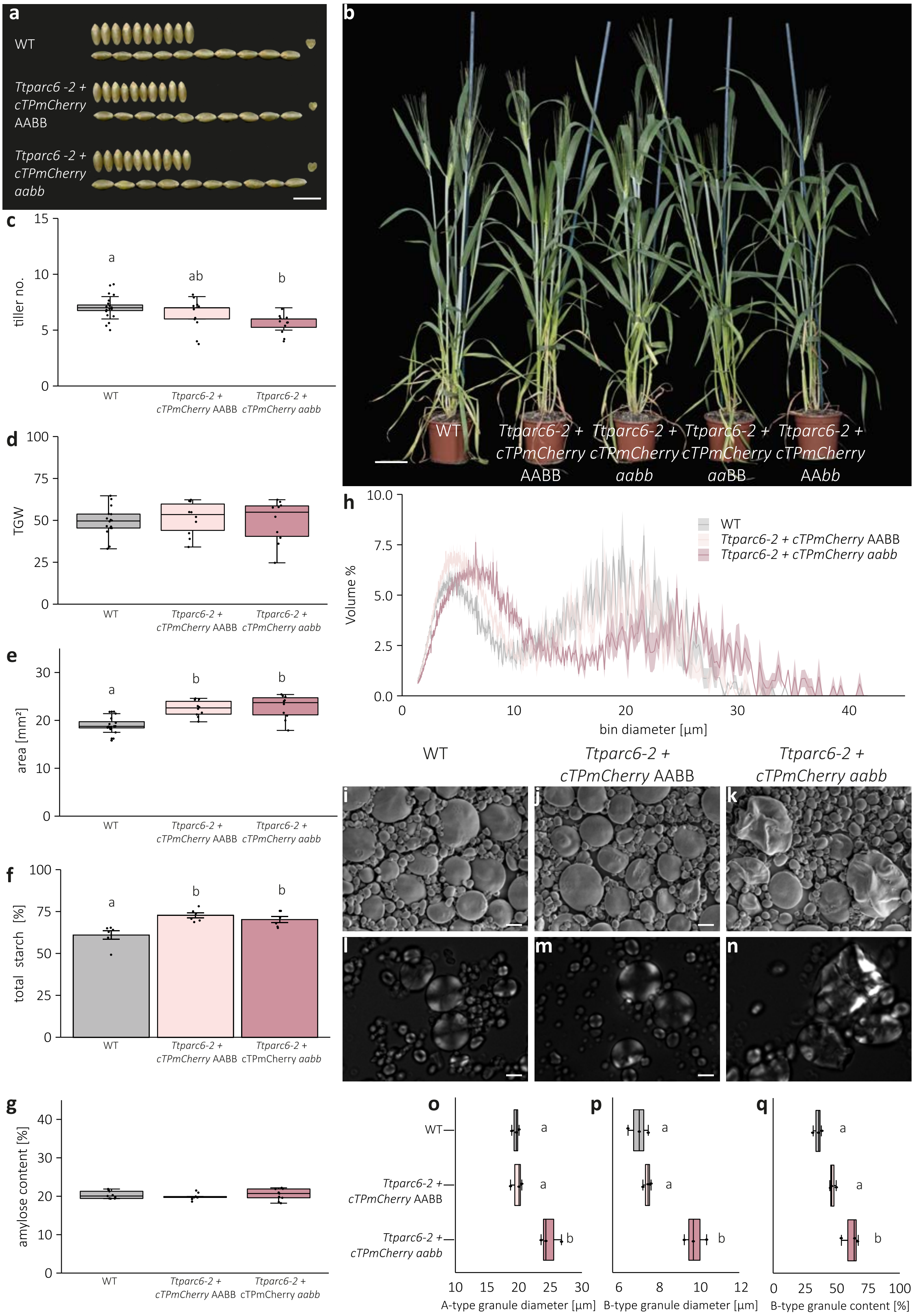


**Figure S6: Plant growth, grain morphology and starch phenotypes of the *Ttparc6* mutant expressing the cTPmCherry amyloplast marker.**
**(a)** Photograph of 10 representative mature grains per genotype. Bar = 1 cm.
**(b)** Photograph of 8-week old transgenic cTPmCherry overexpressing mutant plants and corresponding negative segregants (*Taparc6-2 + cTPmCherry aabb* and *Taparc6-2 + cTPmCherry* AABB) and WT (cv Kronos) plants. Bar = 10 cm.

**(c)** The number of tillers per plant (Tiller no.) of mature plants (n = 10 – 16 per genotype). Significant differences between the lines as determined by Kurskal-Wallis One Way ANOVA on the Ranks all pairwise multiple comparison (Dunn's Method) ($p \leq 0.007$) are represented by different letters.
**(d)** Thousand grain weight (TGW) (in g). Dots represent calculated TGW of individual plants (n = 10 – 16 per genotype). There were no significant differences under a one-way ANOVA.
**(e)** Grain size measured as seed area (in $mm^2$). Dots represent measurements for seeds of 10-16 individual plants per genotype. Significant differences under a one-way ANOVA all pairwise multiple comparison procedures (Tukey's Test) are represented by different letters ($p \leq 0.05$).
**(f)** Total starch content as % (w/w). 3 technical replicates of 2 biological replicates per genotype. Significant differences under a one-way ANOVA and all pairwise multiple comparison procedures (Tukey's Test) are represented by different letters ($p \leq 0.002$).
**(g)** Amylose content [% of total starch]. Dots represent 3 technical replicates of 3 biological replicates. There were no significant differences between the lines determined by Kurskal-Wallis One Way ANOVA on the Ranks (p=0.317).
**(h)** Size distribution plots from Coulter counter analysis. The volume of granules at each diameter relative to the total granule volume was quantified using a Coulter counter. Values represent mean (solid line) ± SEM (shading) of three biological replicates.
**(i-k)** Scanning Electron Microscopy of purified starch granules from mature grain. Bars = 10 µm.
**(l-n)** Polarised light microscopy of purified starch granules from mature grain. Bars = 10 µm.
**(o-q)** Granule size parameters obtained from fitting a log-normal distribution to the B-type granule peak and a normal distribution to the A-type granule peak in the granule size distribution data presented in (h). Three biological replicates were analysed: **(o)** A-type granule diameter (in µm). Significant differences under a one-way ANOVA and all pairwise multiple comparison procedures (Tukey's test) are indicated with different letters ($p \leq 0.02$). **(p)** B-type granule diameter (in µm). Significant differences under a one-way ANOVA and all pairwise multiple comparison procedures (Tukey's test) are indicated with different letters ($p \leq 0.001$). **(q)** B-type granule content by percentage volume. Significant differences under a one-way ANOVA and all pairwise multiple comparison procedures (Tukey's test) are represented with different letters ($p \leq 0.001$).
For all boxplots, the bottom and top of the box represent the lower and upper quartiles respectively, and the band inside the box represents the median. The ends of the whiskers represent values within 1.5x of the interquartile range, whereas values outside are outliers.

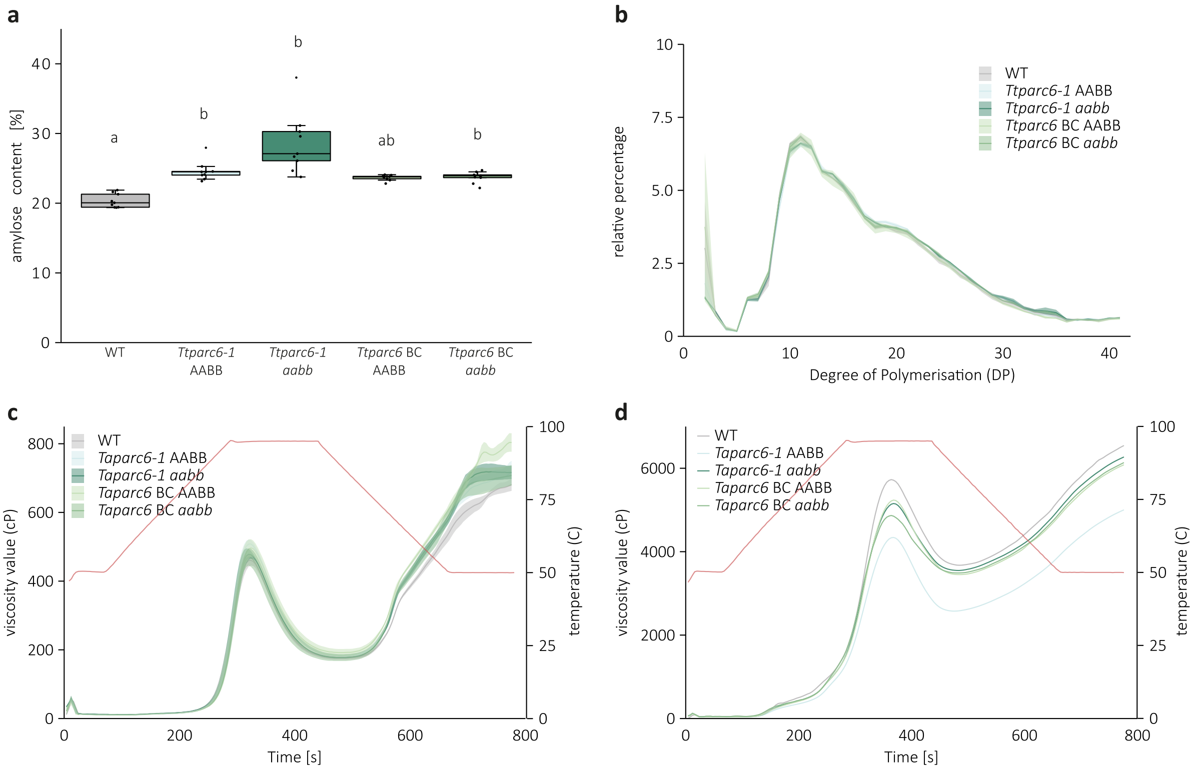


**Figure S7: Analysis of properties of *Tt*PARC6-deficient *Triticum turgidum* starch.**
**(a)** Amylose content [% of total starch] per genotype. Dots represent 3 technical replicates of 3 biological replicates. The bottom and top of the box represent the lower and upper quartiles respectively, and the band inside the box represents the median. The ends of the whiskers represent values within 1.5x of the interquartile range, whereas values outside are outliers. Significant differences between the lines as determined by Kurskal-Wallis One Way ANOVA on the Ranks all pairwise multiple comparison (Tukey's test) ($p \leq 0.001$) are indicated with different letters.
**(b)** Amylopectin chain length distributions. Starch was purified from grains and debranched with isoamylase prior to analysis using HPAEC-PAD. The y-axis represents the relative percentage of chains at each DP. Each line represents the average of three replicates per genotype (each using starch from grains harvested from a separate plant), and the shading represents the SEM.
**(c-d)** Rapid Visco Analyser (RVA) analysis of viscosity during gelatinisation. Analyses were conducted using: **(c)** Purified starch (1.5 g in 25 mL water). Values represent mean (solid line) ± SEM (shading) of three biological replicates, each using starch from grains harvested from a separate plant. The different genotypes produced viscographs that were generally similar, and there was no obvious difference in peak viscosity or the holding strength during cooling. Slight differences between genotypes were observed during cooling (retrogradation) and final viscosity, but these were not consistent for either *Ttparc6* double mutants or wild-type controls. **(d)** Whole flour (5 g in 25 mL water), where flour was produced by pooling a minimum of 3 biological replicates per genotype. The viscographs were more variable among genotypes, and there was no consistent effect that could be attributed to the mutant genotype.

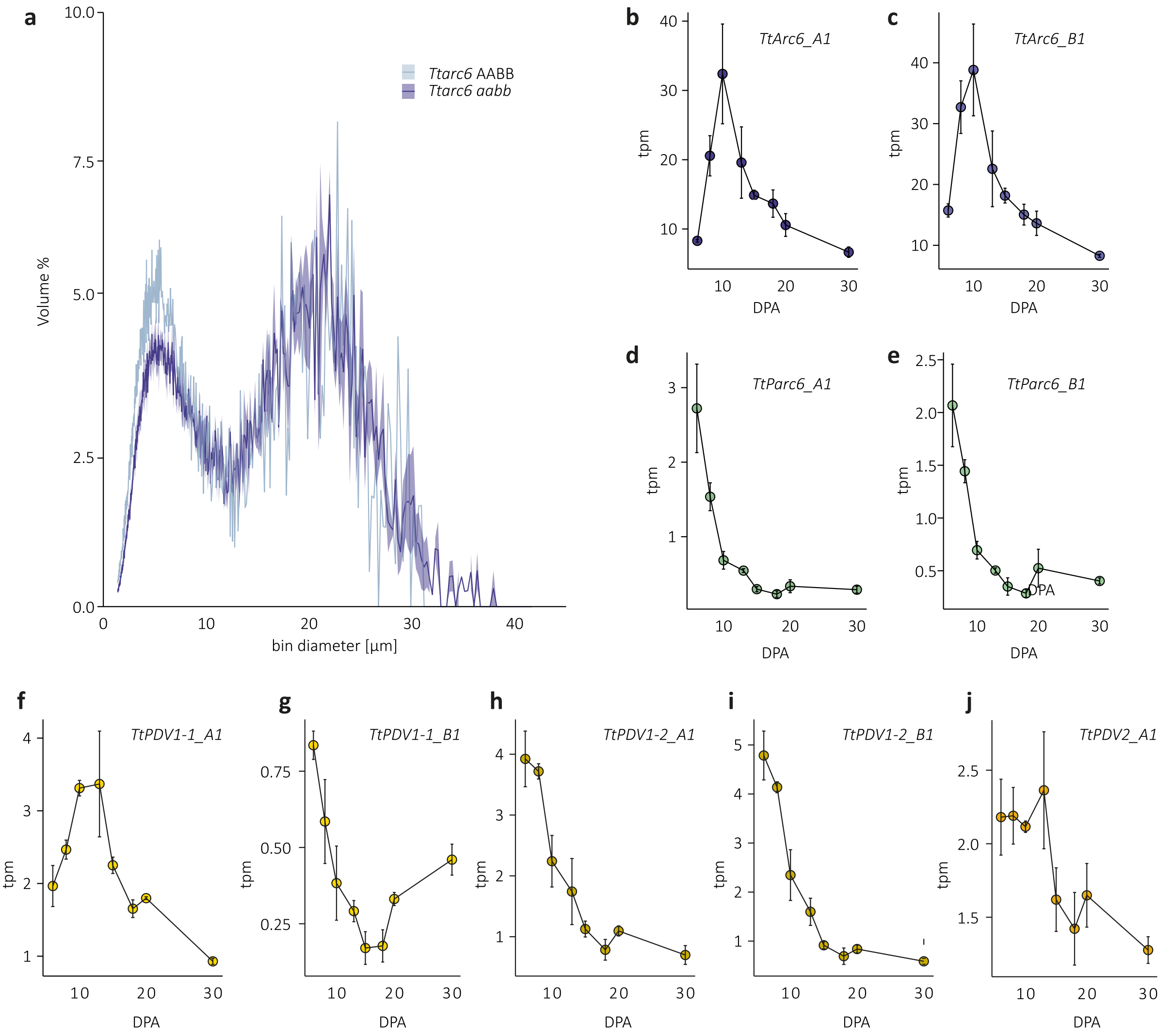


**Figure S8: Size distribution of starch granules of the *Ttarc6* mutant and *TtARC6* and *TtPARC6* expression patterns during endosperm development.**

**(a)** Size distribution of purified starch granules from mature grain. The volume of granules at each diameter relative to the total granule volume was quantified using a Coulter counter. Values represent mean (solid line) ± SEM (shading) of three biological replicates.

**(b-j)** Average TPM values of the *PARC6, ARC6*, *PDV1-1*, *PDV1-2* and *PDV2* homeologs in the durum wheat endosperm. Values are the mean ± SEM of the n = 3 replicates. Values were retrieved from (Chen *et al.*, 2022b).

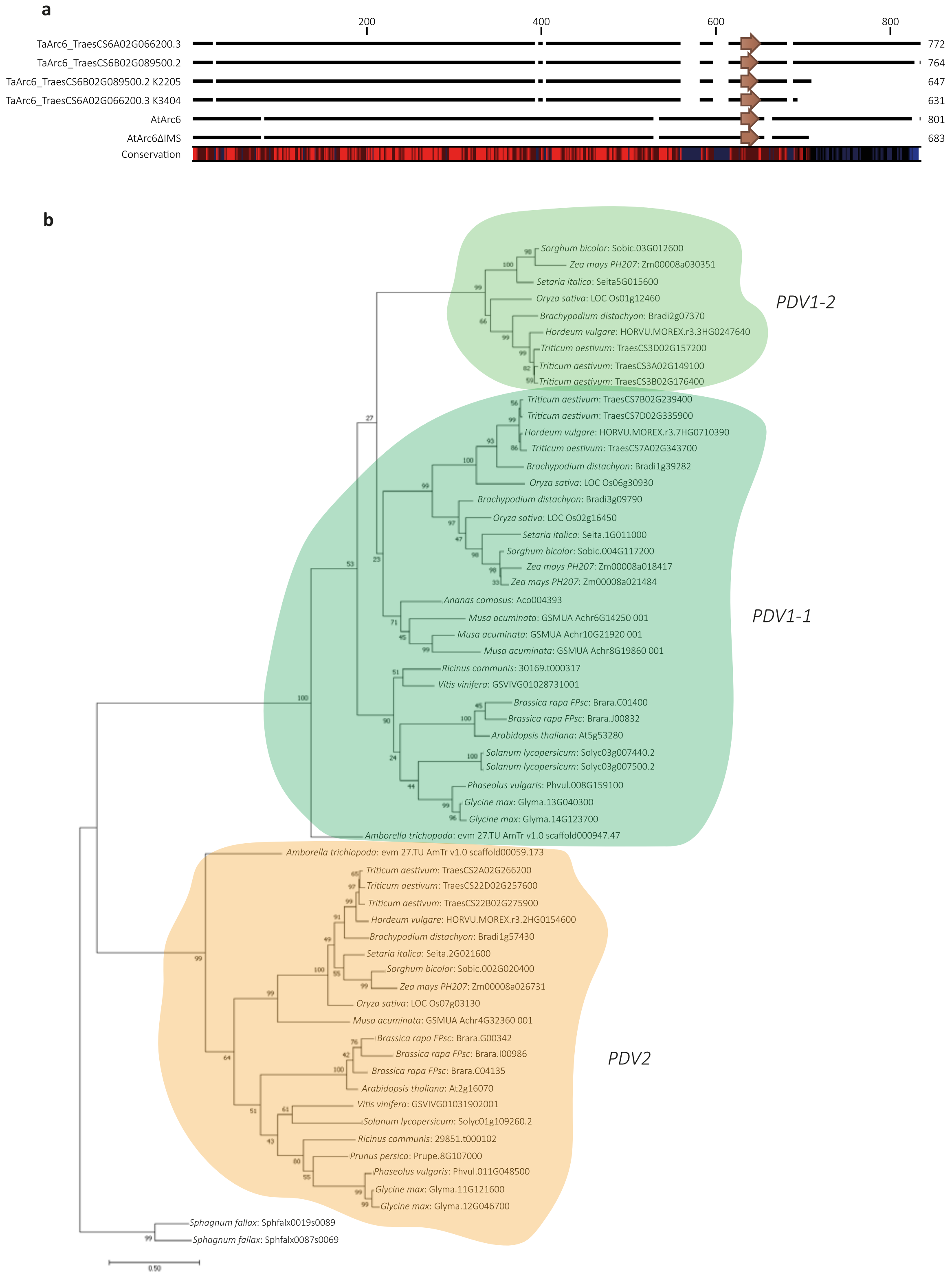


**Figure S9. Molecular phylogenetic analysis of *PDV1* and *PDV2* gene families and *ARC6* mutant protein alignment.**

**(a)** Schematic representation of multiple protein sequence alignment of wheat and Arabidopsis ARC6 protein sequences. Black lines indicate aligned sequence, gaps represent gaps in alignment. Degree of conservation is represented below, where red indicates high conservation and blue indicates low conservation. Brown arrows represent the region encoding the annotated transmembrane domain. Sequences and mutant sequences were retrieved from Ensembl plants and Glynn et al. (2008).

**(b)** Molecular phylogenetic tree of the PDV gene family was constructed from an amino acid alignment using the Maximum Likelihood method based on the JTT matrix-based model. The tree with the highest log likelihood (-20825.25) is shown. The percentage of trees out of 1000 bootstraps in which the associated taxa clustered together is shown next to the branches. The tree is drawn to scale, with branch lengths and scale bar representing the number of substitutions per site. Full experimental procedures can be found in SEP S1.

**Table S1: KASP-markers for *Ttparc6* and *Ttarc6* genotyping.**

| Line | Specificity | VIC/HEX or FAM tail | Genome specific sequence |
|---|---|---|---|
| Kronos1265 | WT | gaaggtcggagtcaacggatt | cgagaagagtccttttgagctctc |
| | Mutant | gaaggtgaccaagttcatgct | cgagaagagtccttttgagctctt |
| | Common | | gcctatccgttgatccctggc |
| Kronos2369 | WT | gaaggtcggagtcaacggatt | tgcaacataccagtgactg |
| | Mutant | gaaggtgaccaagttcatgct | tgcaacataccagtgacta |
| | Common | | cagcttcaaagtaatggaagttccaattcaag a |
| Kronos2205 | WT | gaaggtcggagtcaacggatt | ccttggatttgatactctgc |
| | Mutant | gaaggtgaccaagttcatgct | ccttggatttgatactctgt |
| | Common | | aatttgtttctgaaatcatagtgcc |
| Kronos3404 | WT | gaaggtcggagtcaacggatt | gcatccattctaggaatctg |
| | Mutant | gaaggtgaccaagttcatgct | gcatccattctaggaatcta |
| | Common | | aatttgtttctgaaatcatagtgct |

WT: wild-type,

**Table S2: Codon optimized DNA sequences of *Ta*PARC1-A1, TaARC6-A1, *Ta*PDV1-1-A1, *Ta*PDV1-2-A1 and *Ta*PDV2-A1.**

| codon optimized sequences |
|---|
| >TaParc6_A1_CDS_attL_MluI_codon_optimized_Nbenth<br>ACGCGTCAAATAATGATTTTATTTTGACTGATAGTGACCTGTTCGTTGCAACAAATTGATGAGCAATGCTTTT<br>TTATAATGCCAACTTTGTACAAAAAAGCAGGCTTCACCATGGCTATGCCTACTCCTGCTGCTGCGCTTCTTCA<br>TCCATCTTCTGCTGTGGTAGCTGCTCCTTCTCCTTCTACGTCTTCTTCTGCTAGACGGTCTGCTCCTTCTTCT<br>TCTTCTTCTTCTGCGAGAAGAGGTGGTAATGCTTCTGCTGGTAGAGGTGCCGCTGTTAGACCTAGAGTTGCTG<br>GTGCGGCTGCACCTGTGACAGCTGCTGCTGCTGCTGAAGGTTGTGGTAGACAAGAGCCTCCTGCGGCCCCTGC<br>TGTTGAAATTCCTGTTACTTGTTATCAAATTCTTGGTGTTACTGAAAAGGCTGAAAAGGATGAAATTGTTAAG<br>TCTGCTATTGAACTTAGAAAGTCTGAAATTGAAGATGGTTATACTGAAGAAGTTTCTACTTGTAGACAAGCTC<br>TACTACTTGATGTTAGAGATAAGCTTCTTTTTGAACAAGAATATGCTGGTTCTACTAGAGCTAAGGTTCCTCC<br>TAGATCTTCTCTTCATATTCCTTGGTCTTGGCTTCCTGCTGCTCTTTGTGTTCTTCAAGAAGTTGGTGAAGAA<br>AAGCTTGTTCTTGATATTGGTCAAGCTGCTCTTAGAAGAACTGATTCTAAGCCTTATGCTCATGATGTTCTTC<br>TTGCTATGGCTCTTGCTGAATGTTCTATTGCTAAGGCTTCTTTTGAAAAGTCTAAGGTTTCTCTTGGTTTTGA<br>AGCTCTTGCTAGAGCTCAATATCTTCTTAGAAAAAAACCATCTCTTGAAAAGATGCCTCTTCTTGAACAAATT<br>GAAGAATCTCTTGAAGAACTTGCTCCTGCTTGTACTCTTGAAGTTCTTTCTCTTCCTAGAACTCCTGAAAATT<br>CTGAAAGAAGAAGAGGTGCTATTGCCGCTTTGTGTGAGCTGCTTGGTCAAGGTCTAGATGTTGAATCTTCTTG<br>TAGAGTTCATGATTGGCCTTACTTTTTAGGCCAAGCGATGGACAAATTGCTTGCTACCGAAATTGTTGAACTT<br>CTTTCTTGGGATTCTCTTGCTACTACTAGAAAGAATAAGAAGTCTCTTGAATCTCAATCTCAAAGAGTTGTTG<br>TTGATTTTAATTGTTTTTATAGAGCTATGCTTGCTCATCTTGCTTCTGGTTTTTCTACTAGACAAACTGAACT<br>TATTTCTAAGGCTAAGACTATTTGTGAATGTCTTGTTGCTTCTGAAAATACTGATCTTAAGTTTGAAGAATCT<br>TTTTGTTCTTTTCTTCTTGGTGAAGAATCTGGTGCTACTGTTTTCGAAAAGCTTCAACAACTTCAATCTAATG<br>GTTCTTCTAATTCTAGAAATTATGGTCTTGCTAAGAAGAAGGATTCTTCTGATAAGGTTACTGTTAATCAATC<br>TCTTGAACTTTGGCTTAAGGAAGTTGCTCTTTCTAGATTTGCTGATACTAGAGATTGCCCCCCTTCTCTTGTT<br>AATTTTTTTGCTGCTCCTAAGAGACTTATTTCTACTTCTAAGCAAAAGCTTGGTGCTACTAGAAGAGTTCTTC<br>TTTCTTCTCAAACTCCTTCTTCTGCCTCTACGTGTAATAGAACTTCTGGTCAACAAAATCCTAGACTTAATTC<br>TACTTCTCATCTTGGTGAAGCTGTTAAGCAACTTGCTCCTACTACTCTTGGTGGTCAAGGTTCTACTGATAGA<br>CCTGTTAATGGTCTTTCTACTACTTCTGTTCCTCTTAAGAGAAATCCTGGTTCTCATCCTGTTAGAACTCTTG<br>AATCTTGGGGTCTTACTGGTGATGTTATTGGTAAGATTGCTTATACTGCTGTTCTTGGTCTTGCTCTTTTTGG<br>TACTCTTAAGCTTCTTAGATTTCAATTTGGTAATACTAAGCCTGCTCCTTCTACTAGAGAATCTGCTGCTACT<br>AGTTCTCTCAACGAAGCTTCTCCTTCTGAAGGTTCTTTTATTTCTTCTAGAGTTAGAGAACAATTCGAAAAGC<br>TTTCCAAAATGCTTTGGCTTAATAATAGAGTTCATCTTAGATCTGAAAGATCTGATCTTTCTCCTGGTTCTTC<br>TGATGTTGCTGCTATTGCTAGAAAGGAAAGAATGTCTCTTCAAGAAGCTGAAGCTCTAGTTAAGCAATGGCAA<br>GATATTAAGTCGGAAGCTTTAGGTCCTGATTATGAAATTGATATGCTTTCTGAAGTTCTTGATGGTTCTATGC<br>TTTCTAAGTGGCAAGATCTTGCTCTTTCTGCTAAGGATCAATCTTGTTATTGGAGATTTGTTCTTCTTAATCT<br>TTCTGTTGTTAGAGCTGAAATTCTGCTTGATGAAGCCGGTGATGGTGAAGTTGCTGAAATTAATGCTGTTCTT<br>GAAGAAGCTGCTGAACTTGTTGATGATTCTCAACCTAAGAAGCCCTCTTACTATTCTACTTATGAAGTTCAAT<br>ATTCTCTTAGAAGACAAGATGATGGTTCTTGGAAGATTTGTGAAGCTGCTGTCAGAGACTTATCTGACCCAGC<br>TTTCTTGTACAAAGTTGGCATTATAAGAAAGCATTGCTTATCAATTTGTTGCAACGAACAGGTCACTATCAGT<br>CAAAATAAAATCATTATTTGCACGCGT |
| >TaArc6_A1_CDS_GW_codon_optimized_Nbenth<br>ggggacaagtttgtacaaaaaagcaggcttcATGGAGGGTCTCCACAACCTGCTTGCAAGACCTAATTCTGCA<br>CCTTTAGCATTTTCTCCACCTAGACCAAGACCAAGAAGAAGGCCACCTGTTGCATGTCGAGCAGCTAGCCGCT<br>GGGCCGACCGCCTCTTCGCCGACTTCCATTTATTGCCTACAGCAGCAGCTCCTGAACCACCTGCAGCAGCTCC<br>TGCTGGTGTATCTGCATCACCTTGTGTACCTCTATTTCCAGATGCCGCCGACCGCTCCCTTCCCCTCCAGGTC<br>GACTTCTACAAGGTTCTCGGCGCGGAACCGCATTTCCTCAGCGACGGCGTCAGGCGGGCCTTCGAGGCGAGGG<br>CGGCCAAGCCACCGCAGTACGGCTACAACACAGATACCCTTGTTGGCCGTCGGCAAATACTGCAGCTTGCACA<br>TGATACTCTCACAAACCAGAGCTCCCGCACCGAGTATGACCGCGCGCTCTCTGAGGACCGTGGCATGGCGCTC<br>ACATTGGATGTTGCTTGGGACAAGGTTCCGGGTGTGCTGTGTGCCCTTCAGGAGGCTGGGGAGGCACAGGCAG<br>TGCTCGCAATTGGAGAGCACTTGCTGCAGGACCGCCCGCCCAAGCAGTTCAAACAGGATGTGGTGCTGGCAAT<br>GGCTCTGGCCTATGTGGATCTATCAAGGGACGCAATGGCGGCTAGCCCACCAGACGTAATCCGCTGCTGTGAG<br>GTGCTTGAAAGGGCTCTCAAGCTCTTGCAGGAGGATGGGGCAATCAATCTCGCACCTGATCTGCTTTCACAAA<br>TTGATGAAACTCTGGAGGAGATCACACCTCGTTGTGTTTTGGAGCTTCTTGCCCTTCCTCTTGATGAAAAGCA<br>CCAGAGTAAACGCCAAGAAGGTCTTCGTGGTGTGAGAAACATTTTGTGGAGTGTTGGTAGAGGAGGTATTGCT<br>ACTGTTGGAGGAGGATTTTCGCGTGAAGCCTACATGAATGAGGCCTTTTTGCAGATGACATCAGCGGAGCAGA<br>TGGATTTCTTTTCAAAAACGCCAAATAGCATACCACCTGAATGGTTTGAAATCTATAGTGTGGCACTCGCAAA<br>TGTTGCTCAAGCAATTGTAAGTAAAAGGCCAGAGCTCATCATGGTGGCAGATGATCTTTTCGAACAGCTCCAG<br>AAGTTCAATATAGGTTCTCAATATGCTTATGATAATGAATTGGATCTTGTGTTGGAAAGGGCACTTTGCTCAT |

```
TGCTTGTGGGAGACATTAGCAACTGCAGAATTTGGCTTGCGATTGATAATGAATCCTCACCACATAGAGACCC
CAAAATTGTAGAGTTTATTGTGAACAACTCTAGCATTGACCACCAGGAGAATGATCTTCTTCCAGGCCTGTGT
AAGCTTTTGGAGACTTGGCTTGTCTCAGAGGTTTTCCCTAGGAGCAGAGATACTCGAGGCATGCAGTTTACAC
TTGGAGACTACTACGATGATCCACAAGTTTTAAGCTACCTAGAAATGATGGAAGGTGGTGGTGCTTCTCATTT
GGCTGCTGCTGCTGCTATAGCAAAACTCGGTGCTCAAGCTACAGCTGCGCTTGGTACCGTGAAATCAAGTGCT
ATCCAAGCATTCAACAAGATTTTTCCATTGATAGAACAGCTAGATCGATCAGACATGGAGAATCCTAATGATG
GCCCTGAGGAATCTGTCAATAAATTTGACCAGAAAACTATTATGGGATTTGATATCCGTGATTCCAAAAATGC
TGCCCTGAAGATTGTCTCTGCCAGTGCATTATTTGCTCTGATGACAGTAATAGGCATGAAGTACTTGCCTCGT
AACAAGGTGCTCCCTGCTATTAGAAGCGAGCATAAGTCCATGACAGTTGCTAATGTTGTTGACTCAGTTGATG
ATGATGCACCAGATGAGCCAATACAGATTCCTAGAATGGATGCGAATCTGGCAGAAGGTATTGTTCGCAAGTG
GCAGAGTATCAAATCCAAGGCCTTGGGATCAGATCATTCTTTGGAATCATTGCAAGAGGTTCTTGATGGCAAC
ATGCTGAAGGTATGGAGGGACCGAGCAGCAGAGATCGAGCGCAAAGGCTGGTTCTGGGACTACACGCTGTCCG
ACGTGGCGATTGACAGCATCACCGTCTCCCTGGACGGACGACGGGCGACTGTGGAGGCGACAATTGAGGAGGC
AGGCCAGCTTACCGATGCAACCGACCCCAGGAACAACGATTTGTACGACACTAAGTACACCACCCGGTACGAG
ATGACCTTCACTGGACCAGGAGGGTGGAAGATAACAGAAGGTGCGGTCCTCAAGTCGTCAgacccagctttct
tgtacaaagtggtcccc
```

```
>TaPDV1-1_A1_CDS_GW_codon_optimized_Nbenth
ggggacaagtttgtacaaaaaagcaggcttcaccATGACCGCTTCTTCACCACCACACGATACCAAACAGTTG
GGGACACAAAGGGAGAAGGAAAAGGGTAACCCAAAGAAGAAAGTTATATTGTCAACCGAAGGAGAAGAGAAGA
AGAGCTCTACACGAAGGGCACCAGGTTGCCCTCGAGCTTGCTTACCTTCATCTGCCCCCCATCCCCCTTCCTC
ACTGCAGAGGAGAAGGGCCAACCGTGTTAATGAGATGCGTTGGGACTGGGAGACACCTGCAACAGAAGCAGAG
GCAGAAGCTTTGCAGGAAAGGATATGGGATCTTCATGATAAGCTCTCTCATGCTATTCTCGCCCTGTCCGCCT
GTGCAGGTTTACCGGCTTGCAGGTGCCGTGGCGCACCCAATGGCCACGTAATACTGAAGGGTCAAAGACCCCC
CCAAGGAGGTGGACACGTAGACTTGGCAGCTGCAGCTGCCGCCATGGCTGACGCCAGGGGATTGCACGCCATC
AGGACCGCCCTGGAAGACCTCGAAGGACACCTGCACTTCCTTCGTGACGTTCAATCTCAACAACGAGCTGACC
GTGATGCAGCAATAGCTAGAGTGCAGCAGAGCAGAATACTCCTTGCAGCAAGGTTAGCCGAACACAGGGGCAA
GGGCCATGGAGTGATCGAAGAGGCACTTGGATTTGTCGGTGATGTGCGTGATAAGTCACATTTTGTAAGTCCG
GAGGACGTCTATGGTATGCACTCCCAGAGTGGTGAGGACGAGGAAGATAGGCGTGGGCATGGTTCCAACATGG
TGGTTCGAGTGGTTTCCTGCTCCTTCGCACTCGCCAAAAATATCTTGAGGTTTGAGACCATGGGCAGCGTGCT
GGGTAACGCCACTGTGTTCGCTGTAAGTATGCTGACTTTCCTCCAACTCCACCAGTTAGCTTCAGGAAAGCAA
ATGCCCGCCGTACAATACCGTAGGACAGATAACGTTAGCTTATCCGGCGGCTCTAGAAAGGATACAAAAGGCA
AACACCTCGAGGTGCTTCTCGCCAGGGGTgacccagctttcttgtacaaagtggtcccc
```

```
>TaPDV1-2_A1_CDS_GW_codon_optimized_Nbenth
ggggacaagtttgtacaaaaaagcaggcttcaccATGGAGCCGGAAGAAGCCGAAGCAGTCCTCGAAACAATC
TGGGACTTGCACGACAAGGTTTCTGACGCCATTCATGCTCTGTCTAGGGCCCACTTTCTCAGAGCCGTGCGAA
GAAGAGCAGGTGGCAAACCAGCCGGTGTAGTGCACATAAAGGGGGTCCCTGCCGATGGCGACGAGGCCGCCGA
TCTTAACGCTGTAGCCGAAGAAGCTAGGAGTTTACATGCCATCCGAGCAGCTTTGGAAGACTTGGAGGATCAG
TTCGAGTGTTTCCTCGCTGTTTGTTCCCAACAACAGGCTGAAAGGGATATTGCATTAGCCCGTTTGCAACAAT
CTCATATCATGTTAACAATACGTCTCAAAGAACACCATGGAAACAATCATAAGGTCATAGATGAAGCATTGGA
CTTCGTGCACAATGTGTATCATGACTTTTGGAGTTTCTTATCCGTGAATAAACCTGAAAAAAGTAGGAGCCAC
TCAGGCGCAAACTCTACTAAGGAGACTGGGGACGGGAGCAATTTTTTAGGCTGGATGGTATCTTCATCCTTGG
ATGCCGTTAGAAATTCATTTAACGTCAAAAACTTCGGCGGCTTCCTGGGCAACAGCGCCGTTTTCGCTGTCGG
GATGATTACTATGCTTCAGCTTCACCTGTTAAGTTCAGGTGAGCAGTCATCTTCATGTGGAAAATACAGCTAC
AGAAGAATCAATCGAGATGACAGTTCACAATCACTTGCTGGTCGTTCACGATCCTCCCATCTTGACGTGTTTC
TTGCAAAGAGTgacccagctttcttgtacaaagtggtcccc
```

```
>TaPDV2_A1_CDS_GW_codon_optimized_Nbenth
ggggacaagtttgtacaaaaaagcaggcttcaccATGGAGGGTGAAGAGGAAATTGGGCTCGTCCTCGCCCGA
GCCAGTGATCTCAGGTCACGAATCTCTGCATGTGCAGCCGCAGCTCGACCTCCTCCAAGATTAGGAGCTGGAG
AGGAAGACGATGGAGGCGAAGAGGAAGAAGAAGAGGTGGAGGTGGAATCCCTCGTTGGAATTAACGACGCACT
GGAAAGTTTAGAACGACAATTGGCTTCTCTTCAAGATCTCCAACACCAACAGAGATATGAGAGAGAGACCGTC
CTGAGTCAGATCGACAGGTCTAGGACCAGTTTGCTTAACAAGCTCAAGGAGTACAAGGGCGAAGATTGTGAGG
CAATACATGAAGCAGCCGCCTTTGCCGGGGAAAAGATCGAGAACGACGACGGGCTCATTCTCCCACCGTATAG
TGGTCACGTGACAAATTCATTCGTCCTTGACGACCTGTACCCCGCCAACTACGTTTCCAAACCAAAATGCCTT
CACAACGGCTTACGAAGCGATGGTATGACGGAAGACTCAACTCGAACCAACCGTACTCAAAACCGTATCCCTG
GGACATCCTCAAGAAACTCTAGTGGGGGAATAAGGTCCTTAATAGGTTGGATGGCTAAAACAGCCGTGATGAT
TGTAGGTGCCATCTCCATCATGAAGGCAGCAGGATATGAGCCGACTATAGGGAGAAGCGGCATCAAGCTTGAC
ATAGCAGGGTTACTCGGGAAGGAAGCTGCCGGTGCAAAGGAGCAAGTACCGCCAACACTGCAATGTCCGCCCG
```

| |
|---|
| GGAAAGTGATGGTGCTTGGGGGGGATGGGCGTGCTCACTGTGTCGTCAAAGAAAGAGTAGAGATACCATTCGG<br>GTCTTCCCTTGACGCTCCAAACGCATCATACGGGTTGGGTgacccagctttcttgtacaaagtggtcccc |
| >cTPmCherry_CDS_GW_codon_optimized_Nbenth<br>GGGGACAAGTTTGTACAAAAAAGCAGGCTTCACCATGTCGGCCCTCACTACGTCACAACTTGCAACTTCAGCA<br>ACTGGATTTGGCATTGCTGACAGGTCGGCGCCCTCCTCGCTCTTGCGTCACGGATTCCAAGGCCTTAAGCCCA<br>GATCACCCGCTGGAGGGGACGCCACGTCCCTTAGCGTTACAACAAGCGCACGTGCTACCCCCAAACAGCAACG<br>TTCAGTTCAAAGGGGGTCTCGTCGGTTTCCGTCTGTGGTTGTTTACGCCACGATGGTTAGCAAGGGTGAGGAA<br>GATAACATGGCCATAATCAAAGAATTCATGCGCTTTAAAGTTCATATGGAAGGTTCCGTCAACGGGCACGAGT<br>TCGAGATCGAGGGCGAAGGAGAAGGAAGACCGTACGAAGGAACGCAGACTGCCAAACTCAAAGTTACGAAAGG<br>CGGGCCGTTGCCTTTTGCGTGGGACATACTTTCTCCTCAGTTTATGTACGGGTCCAAGGCGTATGTCAAGCAC<br>CCGGCAGATATACCTGATTACCTTAAACTCTCGTTTCCTGAAGGTTTCAAATGGGAAAGAGTCATGAATTTTG<br>AGGACGGCGGTGTGGTGACGGTGACCCAAGACTCGTCACTTCAAGACGGAGAGTTCATATACAAAGTGAAACT<br>CCGTGGAACAAATTTCCCTTCAGACGGGCCCGTTATGCAAAAAAAGACTATGGGGTGGGAAGCCTCGTCTGAA<br>CGGATGTACCCCGAGGATGGCGCCTTGAAAGGGGAAATTAAGCAGAGGCTTAAACTCAAGGATGGCGGGCACT<br>ACGACGCGGAGGTGAAGACAACCTATAAGGCCAAGAAGCCAGTTCAGTTGCCCGGAGCTTACAATGTGAACAT<br>AAAGCTTGACATAACTTCCCACAATGAAGACTACACCATTGTTGAACAGTACGAAAGGGCGGAGGGTAGACAT<br>TCCACAGGTGGGATGGACGAGCTTTACAAATAAGACCCAGCTTTCTTGTACAAAGTGGTCCCC |
